# Histone bivalency regulates the timing of cerebellar granule cell development

**DOI:** 10.1101/2023.02.02.526881

**Authors:** Kärt Mätlik, Eve-Ellen Govek, Matthew R. Paul, C. David Allis, Mary E. Hatten

## Abstract

Developing neurons undergo a progression of morphological and gene expression changes as they transition from neuronal progenitors to mature, multipolar neurons. Here we use RNA-seq and H3K4me3 and H3K27me3 ChIP-seq to analyze how chromatin modifications control gene expression in a specific type of CNS neuron, the mouse cerebellar granule cell (GC). We find that in proliferating GC progenitors (GCPs), H3K4me3/H3K27me3 bivalency is common at neuronal genes and undergoes dynamic changes that correlate with gene expression during migration and circuit formation. Expressing a fluorescent sensor for bivalent H3K4me3 and H3K27me3 domains revealed subnuclear bivalent foci in proliferating GCPs. Inhibiting H3K27 methyltransferases EZH1 and EZH2 *in vitro* and in organotypic cerebellar slices dramatically altered the expression of bivalent genes and induced the downregulation of migration-related genes and upregulation of synaptic genes, inhibited glial-guided migration, and accelerated terminal differentiation. Thus, histone bivalency is required to regulate the timing of the progression from progenitor cells to mature neurons.

## Introduction

Neuronal development is controlled by gene expression programs that regulate the developmental progression of neuronal progenitor cells as they undergo glial-guided migration and circuit formation. These developmental gene expression programs are thought to be controlled by epigenetic mechanisms, including changes in chromatin accessibility and modifications of histones and DNA (Dixon et al., 2015; Jenuwein and Allis, 2001; Strahl and Allis, 2000). However, even though epigenetic dysregulation is increasingly recognized as an important contributor to neurodevelopmental diseases (Serrano, 2018), the precise mechanisms by which changes in chromatin control gene expression during neurogenesis, glial-guided migration, and maturation in specific types of CNS neurons are largely unknown.

The cerebellar cortex has long provided an important model system for neuronal development, and studies on the cerebellar granule cell (GC) have identified molecular mechanisms that control glial-guided migration (Edmondson and Hatten, 1987; Horn et al., 2018; Solecki et al., 2004; Solecki et al., 2009) and circuit formation (Brickley et al., 1996; van der Heijden and Sillitoe, 2021; Watt et al., 2009; Zhu et al., 2016). Developing cerebellar GCs are an excellent model for addressing the role of chromatin in neuronal development because they are an extremely numerous yet relatively homogenous population of neuronal progenitors that undergo stereotypic and well-characterized developmental steps to become fully functional CNS neurons. GCs are derived from a pool of committed GC progenitors (GCPs) that undergo prolonged clonal expansion in the external granule layer (EGL) of the developing cerebellar cortex (Consalez et al., 2020; Espinosa and Luo, 2008). Following cell cycle exit, immature GCs extend bipolar neurites (parallel fibers) and form a leading process along the radial glia fibers to undergo glial-guided migration. Postmigratory GCs settle in the internal granule layer (IGL) and become multipolar as they extend dendrites and form synapses with ingrowing mossy fiber afferents to form the cerebellar circuitry.

We recently discovered that dramatic changes in the expression of chromatin-modifying genes, including DNA and histone methyltransferases, occur in GCs during the formation of the cerebellar circuitry (Zhu et al., 2016). This finding suggests that regulation of chromatin landscape is critical for GC development. One family of genes that changed significantly encodes the TET enzymes, which control the formation of 5-hydroxymethylcytosine, a DNA modification that we found is required for the transition from migratory to postmigratory GCs through the regulation of ion channel and axon guidance genes (Zhu et al., 2016). These studies from our laboratory and others (Stoyanova et al., 2021) have therefore begun to reveal how changes in DNA modifications impact neuronal development.

Histone post-translational modifications (PTMs) are chemical changes to histones that modulate DNA packaging and chromatin accessibility to transcription factors. Importantly, although histone methylation represents one of the most diverse classes of epigenetic modifications, the roles of individual histone PTMs and their combinations during the development of specific neuron types *in vivo* are only beginning to be understood. Two crucial histone PTMs associated with gene expression are trimethylation of histone 3 at lysine 4 (H3K4me3) and lysine 27 (H3K27me3). H3K4me3 is localized at gene promoters and generally correlates with active gene expression (Bernstein et al., 2005; Santos-Rosa et al., 2002). The regulation of H3K4me3 is critical for development, as loss-of-function in the methyltransferase genes that generate this modification results in embryonic lethality (Cenik and Shilatifard, 2021). By contrast, H3K27me3 is a repressive modification that is enriched on developmentally silenced genes (Barski et al., 2007; Mikkelsen et al., 2007). During embryonic stem cell development, H3K27me3 is thought to repress the expression of selected genes in a lineage-specific manner and thus contribute to lineage decisions during differentiation (Mohn et al., 2008). Loss of H3K27 methyltransferase Ezh2 or other components of the Polycomb Repressive Complex 2 (PRC2) during early neural development induces profound transcriptional dysregulation and impaired neuronal progenitor proliferation, neuronal cell type specification, and differentiation (Feng et al., 2016; Hirabayashi et al., 2009; Zhang et al., 2014), demonstrating that regulation of H3K27me3 is essential at early stages of development.

At most genomic loci, H3K4me3 and H3K27me3 are mutually exclusive, and maintaining their balance is critical for gene expression regulation during development (Cenik and Shilatifard, 2021; Piunti and Shilatifard, 2016). However, during lineage commitment and cell fate determination in pluripotent cells, these two modifications often co-localize at the promoters of developmentally regulated genes (Bernstein et al., 2006; Mikkelsen et al., 2007; Mohn et al., 2008). These so-called bivalent domains are proposed to poise key developmental genes for subsequent lineage-specific activation (Bernstein et al., 2006) or to protect reversibly repressed genes from irreversible silencing (Kumar et al., 2021). Studies in ESC cultures have shown that during developmental progression from ESCs to neuronal progenitor cells, most bivalent promoters reduce to a monovalent status by retaining just one of the two modifications (Mikkelsen et al., 2007). The role of bivalency beyond the regulation of lineage decisions in pluripotent cells is still under investigation and the question remains as to whether bivalent promoters are present in committed cells, including CNS neuronal progenitors, and, if so, whether they are remodeled during neuronal differentiation and maturation. It also remains to be determined how the regulation of H3K27me3 at bivalent and H3K27me3-only genes controls gene expression and developmental progression in immature neurons.

Here, we investigated how the histone modifications H3K4me3 and H3K27me3 regulate glial-guided migration and maturation of a specific type of CNS neuron, the cerebellar GC. We identify H3K4me3/H3K27me3 bivalent domains in proliferating GC progenitors and show that they are enriched at the promoters of neuronal genes. Bivalent domains are dynamically remodeled during GC development, primarily through the regulation of H3K27me3, correlating with developmental gene expression changes. Lastly, we find that perturbing bivalency through the loss of H3K27me3 inhibited glial-guided migration and accelerated neuronal maturation. Together, these data implicate histone bivalency as a specific epigenetic feature controlling the timing of gene expression during cerebellar cortex development and circuit formation.

## RESULTS

### Dynamics of histone PTMs during cerebellar GC development

The genomic localization of histone PTMs is well established in a range of pluripotent and differentiated cells. However, the dynamics of histone PTMs during the development of specific types of CNS neurons *in vivo* are less well studied. To understand how histone PTMs regulate gene expression during the development of a specific type of neuron, we isolated chromatin and RNA from mouse cerebellar GCs at postnatal days 7 (P7), P12, and P21. These ages correspond to neuronal progenitor proliferation, glial-guided migration, and neuronal maturation, respectively (**Fig. 1A**). We then characterized the dynamics of histone PTMs H3K4me3 and H3K27me3, as well as gene expression at these key stages of neuronal development. To determine the genomic localization of H3K4me3 and H3K27me3 across GC development, we performed chromatin immunoprecipitation combined with sequencing (ChIP-seq) from chromatin isolated from proliferating GCPs (as described in (Hatten, 1985)) and postmitotic GC nuclei isolated with fluorescence-activated nuclei sorting (Xu et al., 2018) (**Fig 1B**). To confirm that the isolation methods yielded cell populations enriched for GCs, we evaluated the levels of H3K4me3 at marker genes of different cerebellar cell types identified based on a published scRNA-seq dataset from P60 mouse brain (Saunders et al., 2018). We found high levels of H3K4me3 at *Neurod1*, which is known to be highly expressed in GCs (Behesti et al., 2021), but little or no H3K4me3 signal at genes specific to other cerebellar cell types (**Suppl. Fig. S1A**). These results showed that the isolation methods yielded chromatin from pure populations of GCs.

**Figure 1.**
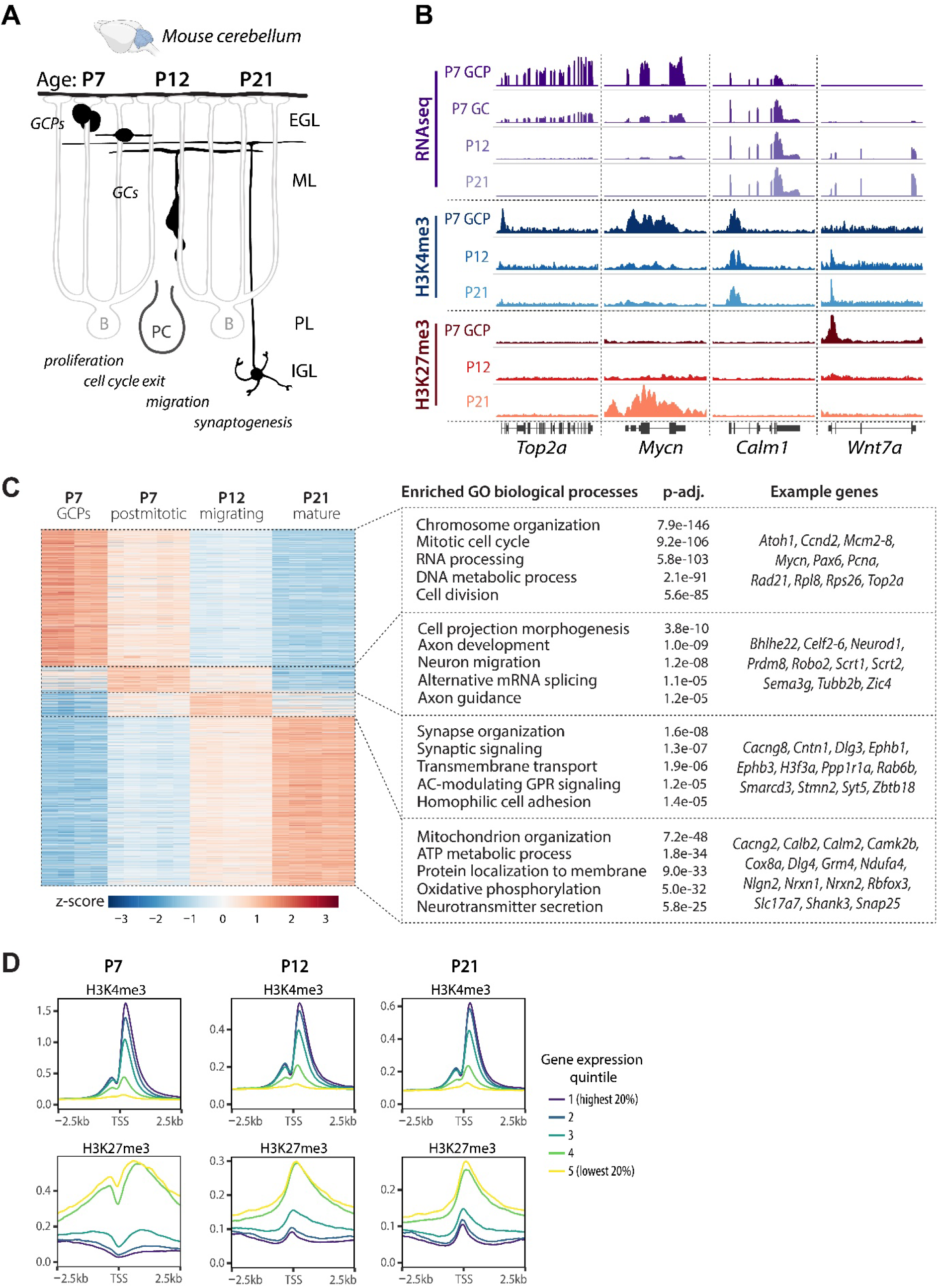
Characterization of gene expression and chromatin landscape in developing mouse cerebellar granule cells. **A** Schematic of postnatal GC development. During late embryogenesis and the first postnatal week, granule cell progenitors (GCPs) undergo clonal expansion in the external granule cell layer (EGL). Upon exiting the cell cycle, immature GCs extend parallel fibers and a leading process and begin glial-guided migration along Bergmann glia fibers. After passing the Purkinje cell layer (PL), the GCs stop migrating and extend dendrites, forming synapses with mossy fiber afferents to form the cerebellar circuitry. IGL, internal granule layer; ML, molecular layer; P, postnatal day. **B** Genome browser view of representative genes undergoing gene expression and histone modification changes during GC development. Representative ChIP-seq and TRAP RNA-seq samples are shown. H3K4me3 (n=2-3 samples/group) and H3K27me3 (n=4-5 samples/group) ChIP-seq was performed on chromatin isolated from GCPs (P7) or from GC nuclei sorted using FANS (P12 and P21). RNA-seq was performed on TRAP RNA isolated from P5-P7 *Tg(Atoh1-Egfp-L10a)* GCPs (P7-GCP, n=4 samples/group), or from the cerebellar lysates of *Tg(Neurod1-Egfp-L10a)* mice at P7 (P7-GC), P12 and P21 (n=5 mice/group). **C** Left: Heatmap depicting differentially expressed (DE) genes in developing GCs. DE genes (p-adj < 0.01, log_2_FC≥1) were identified by pairwise comparisons between groups using DESeq2 and sorted by the highest expressed genes at each age. Right: Gene Ontology (GO) analysis of the highest expressed genes at each developmental time point, identified using clusterProfiler. The GO biological process categories were sorted by the adjusted P-value and the top 5 enriched non-redundant categories are shown for each age. **D** Relative signal of the histone modifications H3K4me3 and H3K27me3 around transcription start sites (TSS) at P7, P12 and P21, grouped into quintiles based on gene expression levels.

To relate histone PTM changes to changes in gene expression, we used the translating ribosome affinity purification (TRAP, (Doyle et al., 2008; Heiman et al., 2008)) and performed RNA-seq on translating mRNA isolated from *Tg(Atoh1-Egfp-L10a)* and *Tg(Neurod1-Egfp-L10a)* mice to characterize gene expression in proliferating and postmitotic GCs, respectively (**Suppl. Fig. S1A-C**). At each developmental time point, known GC-specific genes were enriched in the TRAP samples compared to the input, whereas marker genes of other cerebellar cell types were depleted (**Suppl. Fig. S1A, D**). In addition, the expression of marker genes of other cerebellar cells was significantly lower than the expression of GC-specific genes in TRAP RNA (**Suppl. Fig. S1E**), demonstrating that the TRAP methodology yielded GC-enriched RNA.

We then determined the gene expression patterns of developmentally regulated genes in GCs at each developmental stage. To that end, we performed pairwise differential gene expression analysis using DESeq2. Genes with a log2FC≥1 and p-adj<0.01 in any pairwise comparisons were considered to be differentially expressed. For each gene, we determined the developmental stage where this gene was expressed at the highest level and subsequently performed Gene Ontology (GO) enrichment analysis to identify the biological processes enriched at each developmental stage (**Fig. 1C**). At P7, the enriched pathways in proliferating GCPs were associated with the cell cycle, whereas newly differentiated, postmitotic GCs predominantly expressed genes involved in axon extension and axon guidance. In migrating GCs at P12, genes expressed at the highest levels were related to neuronal differentiation and synaptic signaling, whereas pathways involved in mitochondrial function and neurotransmitter secretion reached their highest expression in mature GCs at P21. These data were consistent with a previously published microarray dataset from our laboratory (Zhu et al., 2016). We next performed a genome-wide comparison of ChIP-seq and RNA-seq data and found that the levels of H3K4me3 and H3K27me3 at the promoter correlated with gene expression, with the H3K4me3 signal being highest at highly expressed genes and H3K27me3 at repressed genes (**Fig. 1D**).

### GCPs exhibit developmentally regulated H3K4me3/H3K27me3 bivalency

In addition to identifying genes with H3K4me3 and H3K27me3 marks at specific developmental stages, we discovered that these modifications often fully or partially overlapped, forming bivalent domains. Bivalent regions almost exclusively localized to promoters (**Fig. 2A**). We determined H3K4me3 and H3K27me3 profiles at H3K4me3-only, bivalent, and H3K27me3-only genes at P7, and found that both H3K4me3 and H3K27me3 signals were on average higher at monovalent than bivalent loci (**Fig. 2B**). We noticed that genes with bivalent promoters were often bivalent in GCPs and resolved to monovalent in mature GCs (**Fig. 2C**). For example, genes encoding the cell cycle protein cyclin D1 (*Ccnd1*) and the voltage-gated potassium channel subunit *Kcna1* were both bivalent at P7, but during migration and maturation *Ccnd1* retained H3K27me3 with a corresponding decrease in gene expression, while *Kcna1* retained H3K4me3 with a corresponding increase in gene expression (**Fig. 2C**). We also identified genes that underwent a more dynamic change in bivalency, like *Cxcl12*, for example, which encodes a chemokine involved in GC glial-guided migration (Ozawa et al., 2016). We found that *Cxcl12* was bivalent at P7 and P21, but during glial-guided migration at P12 H3K27me3 was transiently removed, correlating with a peak in Cxcl12 expression in migrating cells (**Fig. 2C**).

**Figure 2.**
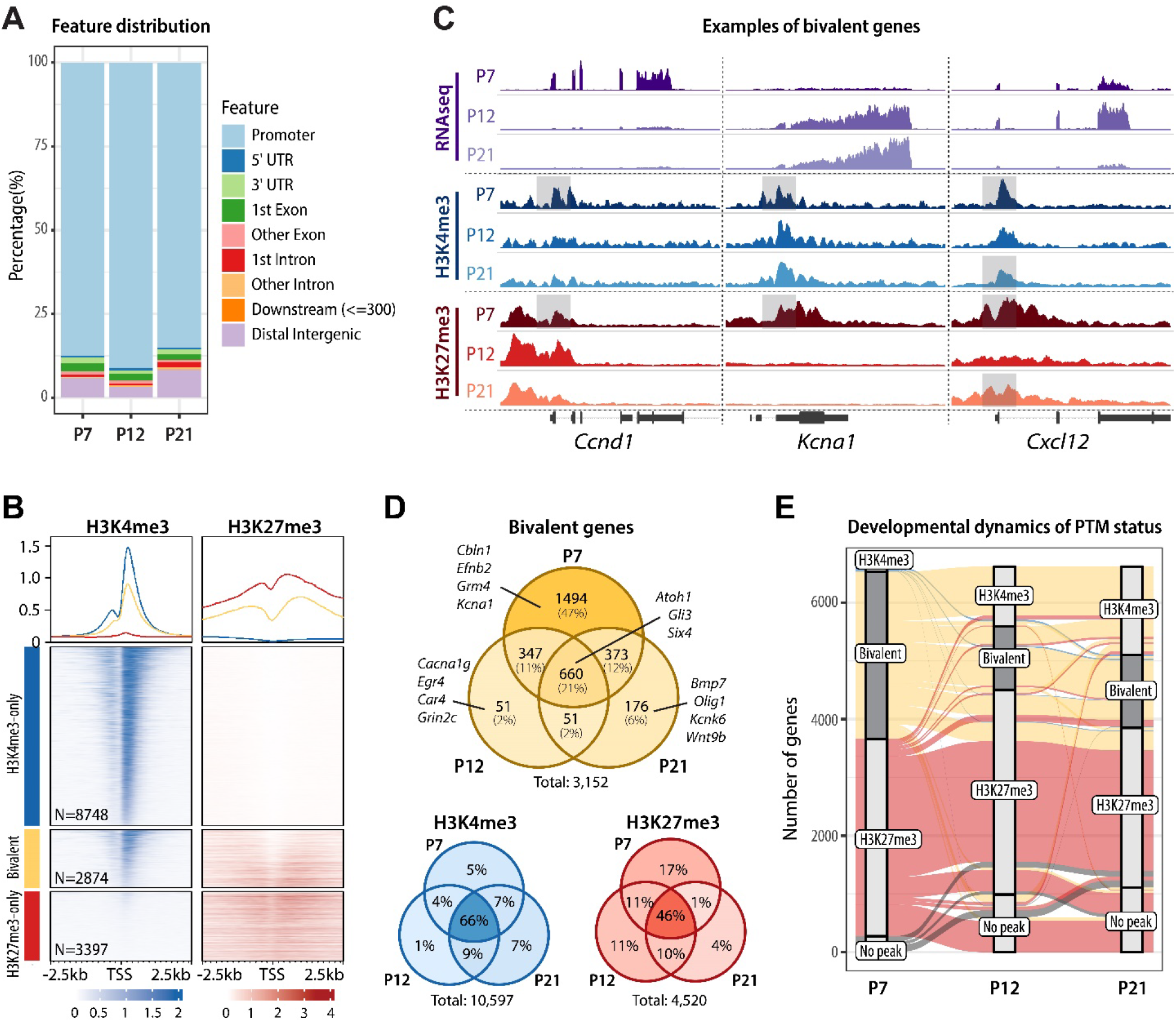
H3K4me3/H3K27me3 bivalent promoters are prevalent in proliferating GC progenitors. **A** Genomic distribution of bivalent peaks. Bivalent peaks were defined as regions where H3K4me3 and H3K27me3 peaks overlapped. The genomic distribution of the identified overlapping regions, defined as bivalent, was determined using the ChIPseeker package. **B** Enrichment map depicting ChIPseq normalized reads in P7 GCPs, centered at TSS ± 2.5 kb, sorted by H3K4me3 signal, and grouped by TSS status (H3K4me3-only, bivalent, or H3K27me3-only). **C** Genome browser view or representative bivalent genes. Shaded areas denote the bivalent regions around the TSS. Note that *Ccnd1* and *Kcna1* are both bivalent at P7 but during migration and maturation *Ccnd1* retains H3K27me3 and *Kcna1* retains H3K4me3 with a corresponding change in gene expression. *Cxcl12* is bivalent at P7 and P21, while H3K27me3 is transiently removed at P12, corresponding to a peak in its expression at P12. **D** Venn diagrams depicting the number of bivalent, H3K4me3-only, and H3K27me3-only genes across GC development. Examples of bivalent genes are shown. Note that bivalent genes are particularly common in P7 GCPs, while the majority of H3K4me3-only and H3K27me3-only genes are stable between P7 and P21. **E** Alluvial plot showing the dynamics of histone post-translational modifications between P7 and P21. Bars represent PTM statuses at promoters and lines indicate the changes in PTMs over development. Only genes that are either bivalent or H3K27me3-only at any age are shown. Bivalent genes are highlighted. Groups that contain fewer than 0.5% of included genes are omitted for simplicity. Note that a major fraction of P7 bivalent genes become H3K4me3-only by P12 and remain so through P21, whereas most P7 H3K27me3-only genes remain H3K27me3-only until P21.

Across all developmental stages, we identified a total of more than 3000 bivalent genes (**Fig. 2D, Supplementary Table S1**), demonstrating that bivalency is common in a committed CNS neuron progenitor population. The number of bivalent genes was high in progenitor cells and markedly lower in migrating and mature neurons (**Fig. 2D**). Over development, approximately 50% of genes bivalent at P7 resolved to H3K4me3-only by P21, demonstrating that H3K27me3 was actively removed from bivalent loci during GC differentiation and maturation (**Fig. 2E**). In contrast, the methylation pattern of genes that only had H3K4me3 or H3K27me3 was relatively stable as development progressed (**Fig. 2D-E**, **Suppl. Fig. S2A**). Moreover, regulation of H3K27me3 was more dynamic at bivalent loci than at H3K27me3-only loci: most (63%) of P7 H3K27me3-only peaks maintained their H3K27me3-only status throughout GC development, whereas H3K27me3 was removed from 51% of bivalent promoters by P21 (**Fig. 2E**). These results suggest that monovalent genes retain relatively stable methylation status, while PTMs at bivalent genes are dynamically regulated during development.

### Bivalent domains exist in individual GCPs

To address the question as to whether bivalent domains exist in individual GCs or are an averaged feature of the GC population, we used a recently developed multivalent chromatin sensor (cMAP3) that selectively binds H3K4me3/H3K27me3 bivalent nucleosomes (**Fig. 3A**, upper panel, (Delachat et al., 2018; Hoffman et al., 2020)). This genetically encoded sensor contains a fluorescent reporter fused to H3K27me3-binding Polycomb chromodomain (PCD) at the N-terminus and H3K4me3-binding plant homeodomain (PHD) derived from TFIID subunit 3 (TAF3) at the C-terminus. The binding of both reader domains to bivalent nucleosomes leads to the stabilization of fluorescent signal and the appearance of characteristic fluorescent foci at bivalent domains (Delachat et al., 2018; Hoffman et al., 2020). The plasmid also expresses tdTomato from an internal ribosome entry site, allowing us to simultaneously assess GC morphology.

**Figure 3.**
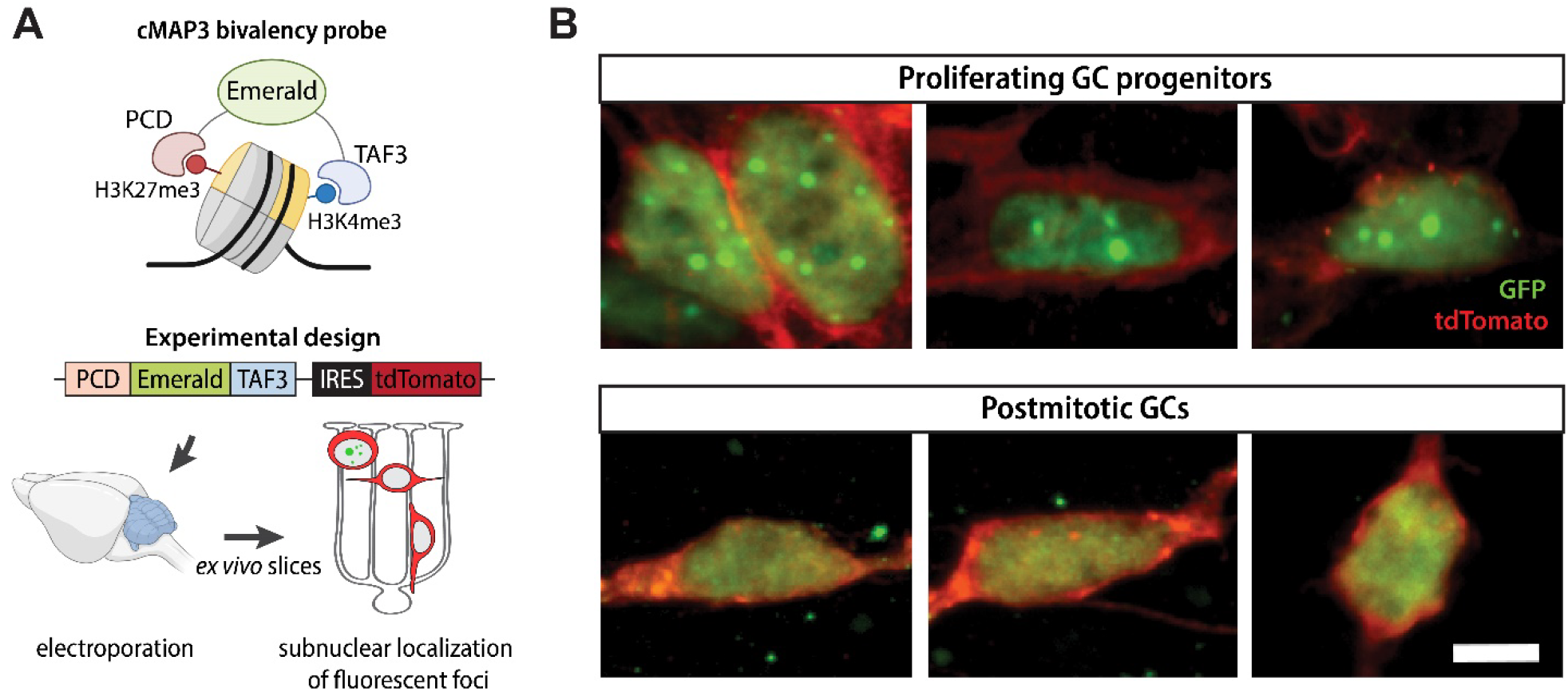
Identification of bivalent domains *ex vivo*. **A** Schematic of the cMAP3 bivalency probe used to detect truly bivalent domains *ex vivo*. The probe consists of a fluorophore (Emerald) fused with H3K27me3 and H3K4me3 reader domains. The plasmid co-expresses tdTomato under IRES for assessing GC morphology. **B** Representative images of GCs electroporated with the cMAP3 bivalency probe, showing the subnuclear localization of bivalent domains. Proliferating GCPs, imaged 6-12h after electroporation, contain multiple fluorescent foci per nucleus, indicative of bivalent domains. Postmitotic GCs, imaged 48-72h after electroporation, contain no foci. Scale bar, 5 μm.

To detect bivalent chromatin in GCs at defined stages of development, we electroporated cerebella from P8 mice with a plasmid co-expressing the cMAP3 construct and the tdTomato reporter (**Fig. 3A**, lower panel), generated *ex vivo* organotypic slices, and imaged cMAP3-expressing cells after 6-72h. As we previously showed, electroporation of the cerebella at this age results in specific targeting of GCPs that reside in the EGL. Importantly, GCs undergoing specific stages of development have the same morphology and laminar position in the *ex vivo* slices as they do *in vivo* (**Fig. 1A**). In addition, GC development in slices follows the same schedule as *in vivo*. Therefore, the *ex vivo* slice culture system allows us to selectively target GCPs and identify GCs at specific stages of development using their morphology and location in the tissue (Govek et al., 2018; Solecki et al., 2004; Zhu et al., 2016).

We imaged cMAP3-expressing GCs at various time points after electroporation. The developmental stage of proliferating GCPs and postmitotic GCs was inferred based on cell morphology, evaluated by tdTomato fluorescence. Cycling GCPs in the outer EGL exhibit a range of complex, dynamic morphologies and can have extensive, elaborate protrusions (Hanzel et al., 2019), while newly differentiated GCs in the lower EGL extend bipolar parallel fiber processes. GCs then extend a leading process in the direction of migration, while trailing a T-shaped axon behind the soma, and they become multipolar as they mature. We found that proliferating GCPs imaged 6-12h after electroporation displayed distinct fluorescent subnuclear foci, whereas postmitotic GCs imaged 48-72h after electroporation exhibited a diffuse fluorescence pattern without detectable subnuclear foci (**Fig. 3B**). These results show that individual GCPs contain bivalent domains and support our observation that bivalency is more prevalent in proliferating progenitor cells than in postmitotic cells (**Fig. 2C-D**).

### Bivalency is enriched at developmentally regulated neuronal genes

Having confirmed that bivalent domains exist in individual GCs, we asked which biological processes were enriched among bivalent genes. Toward that end, we performed Gene Ontology (GO) analysis of all genes that were bivalent between P7-P21 and found that bivalency is common at genes associated with key functions during neuronal development, including neurogenesis, migration, and synaptic function (**Fig. 4A**). We also determined the enriched categories of bivalent and H3K27me3-only genes separately for each developmental stage. Interestingly, we found that the enriched categories of both bivalent and H3K27me3-only genes were remarkably stable between P7 and P21, with bivalent genes being associated with neuronal development and H3K27me3-only genes being associated with earlier developmental processes, such as organ morphogenesis and pattern specification (**Suppl. Fig. S2B**). This suggested that genes marked with both H3K4me3 and H3K27me3 are functionally distinct from genes with only H3K27me3, and further implied that the regulation of bivalency at neuronal genes could be associated with gene expression regulation.

**Figure 4.**
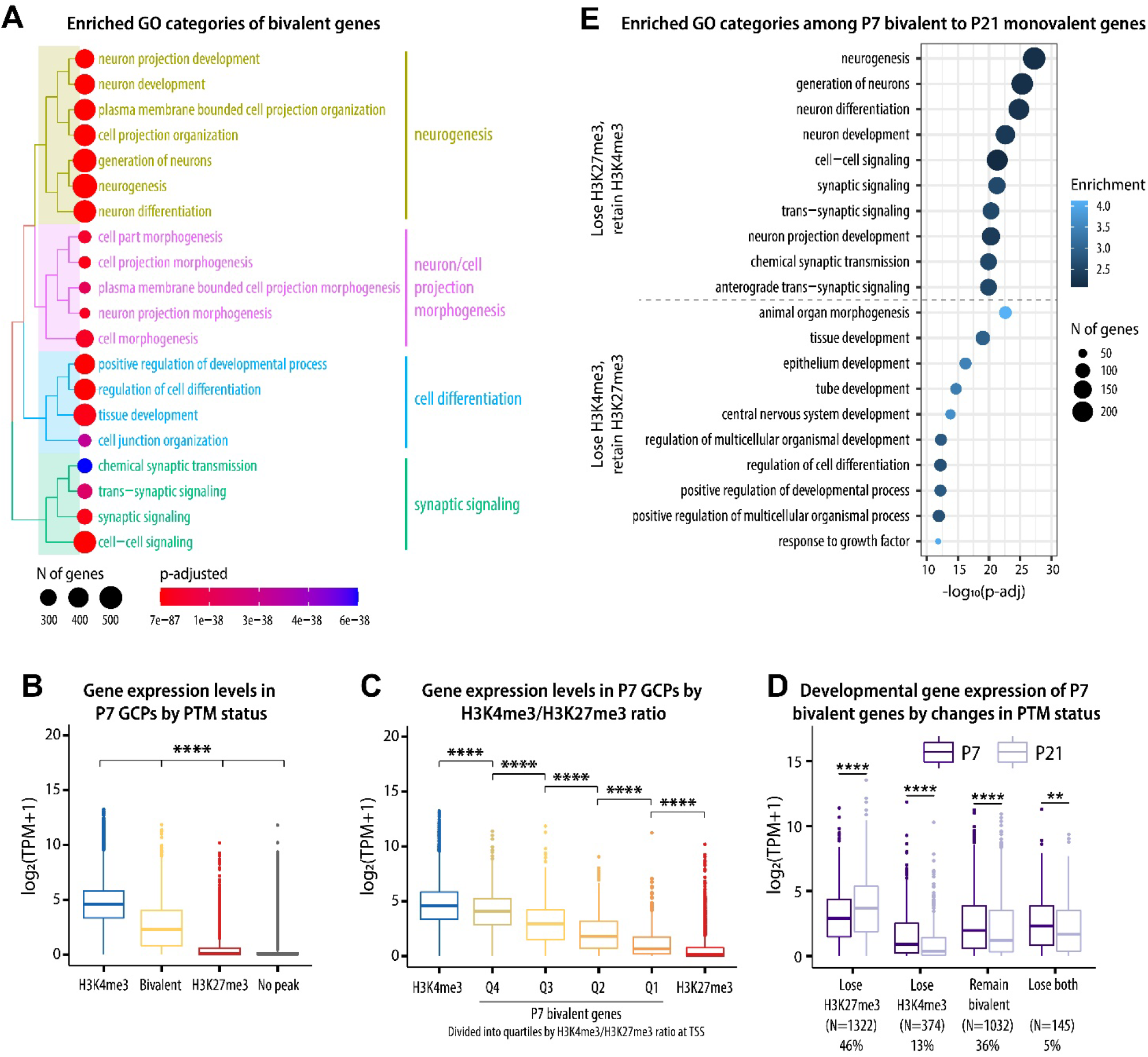
H3K4me3/H3K27me3 bivalency predicts gene expression levels in GCs. **A** Enriched GO (biological process) categories of bivalent genes, identified using the clusterProfiler package. Top 20 identified categories were plotted using the treeplot function to visualize the relationship between the categories. **B**. At P7, bivalent genes are expressed at lower levels than H3K4me3-only genes but at higher levels than H3K27me3 or no-peak genes. Kruskal-Wallis one-way ANOVA, followed by Dunn’s *post hoc* test. **** p < 0.0001 between all comparisons. **C** The expression of bivalent genes correlates with H3K4me3/H3K27me3 ratio at the TSS. One-way ANOVA, followed by Tukey HSD *post hoc* test. Correction for multiple comparisons was performed considering all comparisons but only significances between adjacent groups are shown. **** p < 0.0001. **D** Changes in the PTM status of bivalent genes correlate with gene expression across GC development. Pairwise t-test, adjusted for multiple comparisons using the BH method. ** p < 0.01, **** p < 0.0001. **E** Enriched GO (biological process) categories among genes that change from bivalent to monovalent between P7 and P21, identified with clusterProfile.

Based on our observation that bivalent genes are especially enriched among neuronal genes, we next asked how developmental changes in bivalency correlate with gene expression during GC development. We analyzed the expression of genes with different PTM statuses at promoters (H3K4me3-only, bivalent, H3K27me3-only, and no-peak), and found that genes with bivalent promoters were expressed at lower levels compared to genes with H3K4me3 alone, but at higher levels than genes with only H3K27me3 or genes without either of the peaks (**Fig. 4B**, **Suppl. Fig. S3A**). Moreover, the ratio of H3K4me3 to H3K27me3 at bivalent genes was positively correlated with gene expression (**Fig. 4C**), showing that the balance between H3K4me3 and H3K27me3 signals could be relevant for the regulation of gene expression levels.

We then analyzed the relationship between histone PTM changes and gene expression during GC development and found that changes in chromatin correlated with changes in gene expression levels (**Fig. 4D, Suppl. Fig. S3B**). Interestingly, most genes bivalent in GCPs were differentially expressed during development (44% upregulated and 38% downregulated by P21) (**Suppl. Fig. S3C**). GO analysis revealed that the genes that lost H3K27me3 and retained H3K4me3 were involved with neuronal differentiation and maturation, whereas the genes that lost H3K4me3 and retained H3K27me3 were associated with functional categories related to earlier developmental processes, such as tissue development and morphogenesis (**Fig. 4E**). Therefore, these results suggested that the regulation of H3K4me3 and H3K27me3 levels at bivalent domains has the potential to control and fine-tune the expression of genes required for GC maturation, and that both activation and repression of bivalent genes are required for GC maturation.

### Transcription factors that determine GC identity are bivalent in developing GCs

After establishing that dynamics of bivalency correlated with gene expression, we asked if bivalency could directly control the expression of transcription factors (TFs) that regulate GC developmental progression. We identified a total of 107 bivalent TFs with an average TPM>5. Bivalent TFs included almost all genes known to be essential for specifying GC identity and for GC differentiation, including *Atoh1, Mycn, Pax6, Gli1* and *Neurod1* (**Fig. 5A**). The exceptions are the *Zic* genes that were not bivalent at any age. Moreover, several TFs of the AP-1 complex that regulates activity-dependent gene expression were bivalent in developing GCs (**Fig. 5A**).

**Figure 5.**
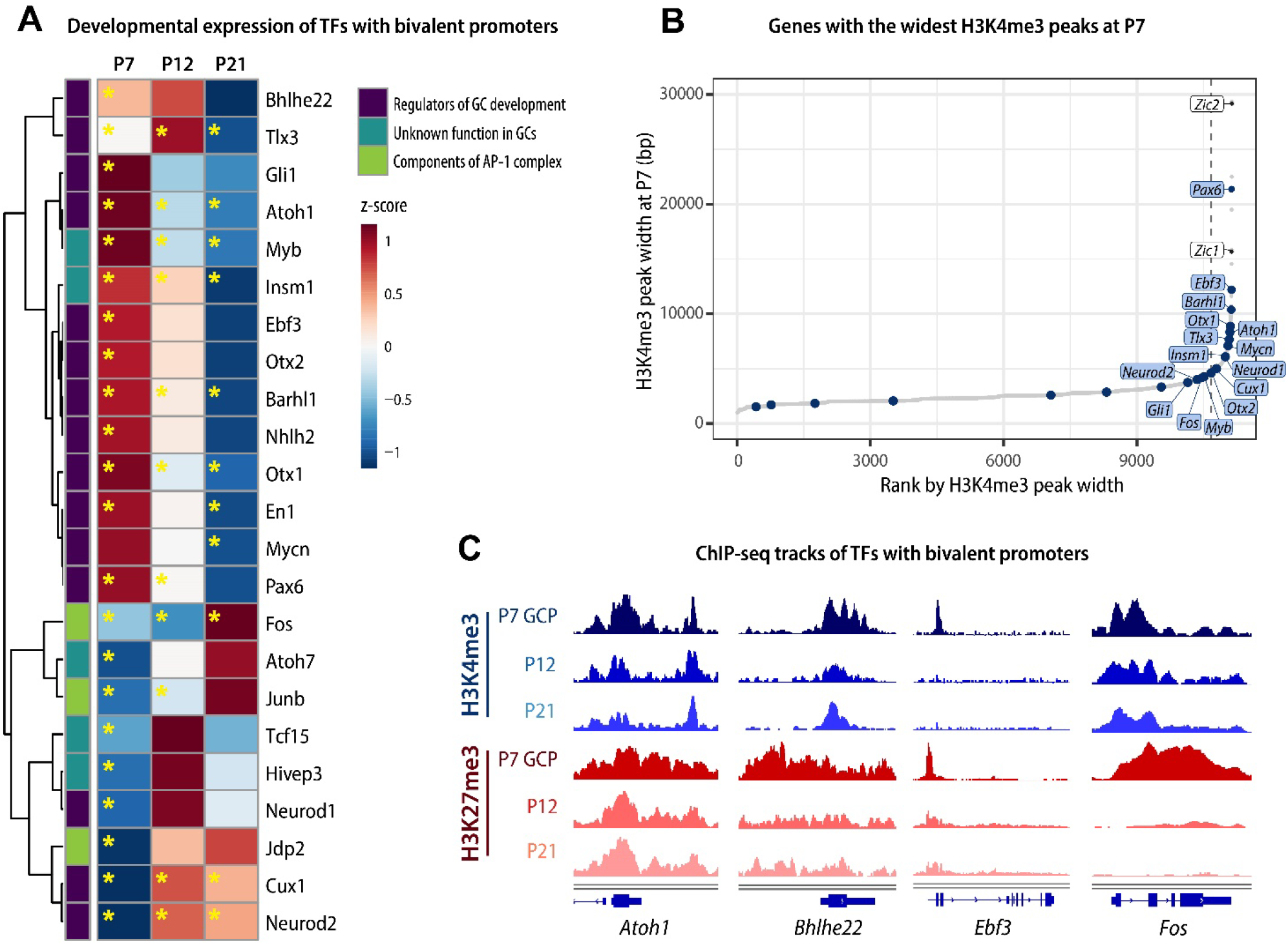
Transcription factors that establish GC identity are bivalent. **A** Heatmap depicting the developmental gene expression dynamics of selected transcription factors with bivalent promoters. Yellow asterisks denote the time points when the genes are bivalent. **B** P7 H3K4me3 peaks ranked by width in base pairs. The dashed line denotes the cut-off point between ‘broad’ and ‘typical’ peaks, determined using the elbow method. Peaks annotated to the promoters of bivalent TF genes are shown in blue. **C** Integrative Genome Browser view of selected transcription factors with bivalent promoters.

We next identified the genes with the broadest H3K4me3 domains, which is an epigenetic signature of cell identity genes (Benayoun et al., 2014). We found that many of the these transcription factor genes had the broadest H3K4me3 peaks (**Fig. 5B-C**), consistent with the known importance of these genes in GC development. Therefore, the TFs that specify and maintain GC identity undergo dynamic changes in their bivalency status during GC neurogenesis, glial-guided migration, and maturation.

### Perturbations in bivalency inhibit glial-guided migration and accelerate terminal differentiation

We next asked whether perturbations in bivalency impact the developmental progression of immature GCs. To address this, we targeted H3K27me3, because our results suggested that H3K27me3 at bivalent loci is more dynamically regulated than H3K4me3 (**Fig. 2D**, **Suppl. Fig. S2A**). To reduce the levels of H3K27me3, we focused on the methyltransferases that generate this modification. H3K27 is methylated by the Polycomb Repressive Complex 2 (PRC2) that contains Enhancer of Zeste 1 (EZH1) or EZH2 as the catalytic subunit. Both EZH1 and EZH2 are expressed throughout GC development, with EZH2 being highly expressed in proliferating GCPs and EZH1 in postmitotic cells (**Fig. 6A**). Importantly, even though EZH2 is the major H3K27 methyltransferase in proliferating cells, results from one study have suggested that bivalent genes are core targets of EZH1 (Aoyama et al., 2018). Therefore, to ensure that our approach to reducing H3K27me3 impacts H3K27me3 at bivalent genes, we used a small molecule inhibitor UNC1999 that inhibits both EZH1 and EZH2 (Konze et al., 2013).

**Figure 6.**
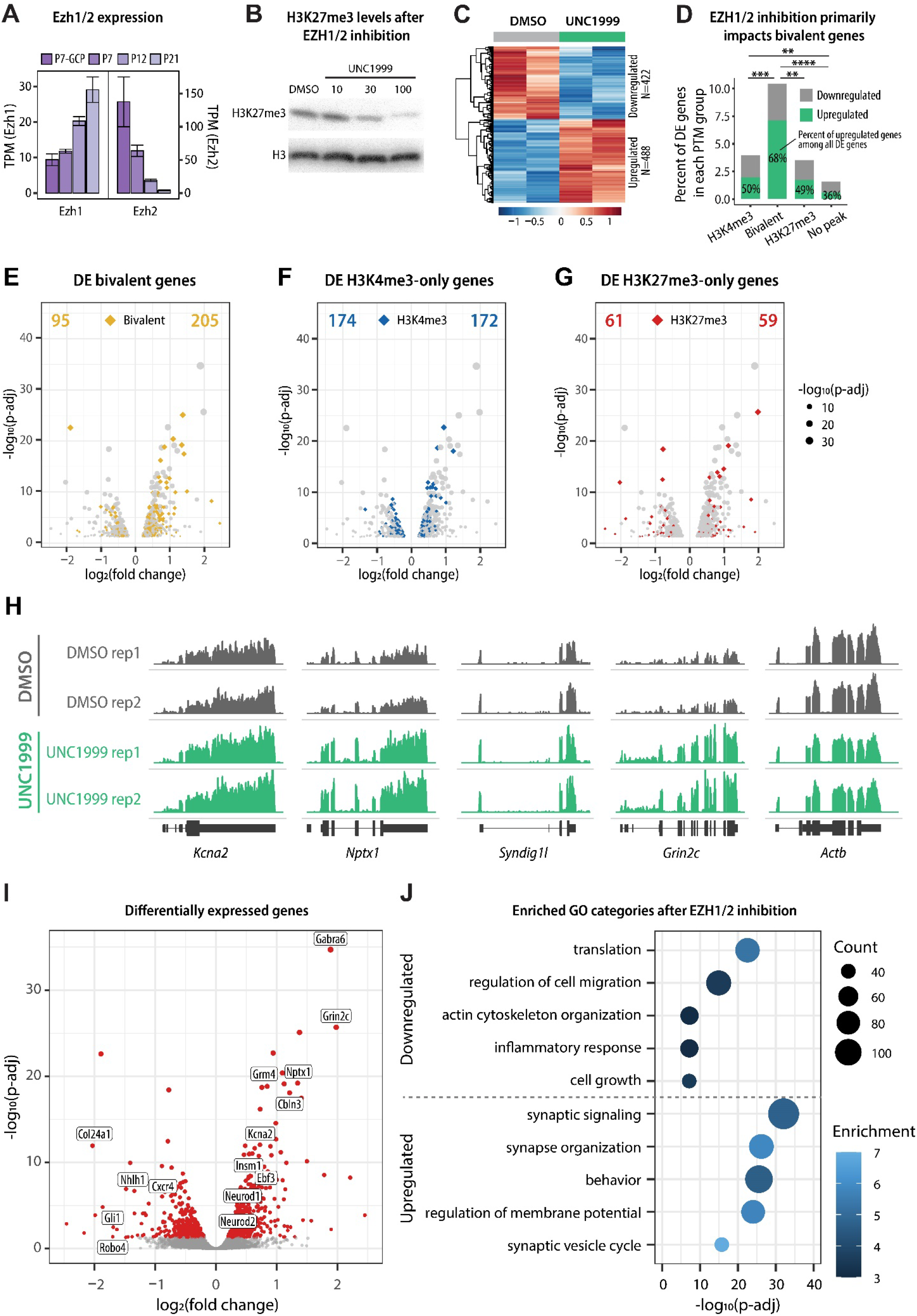
The effect of EZH1/2 inhibition on gene expression in cultured GCs. **A** Developmental expression of H3K27 methyltransferases Ezh1 and Ezh2, based on TRAP RNAseq (n=4-5/group). **B** Immunoblot detection of H3K27me3 levels in GCs cultured for 5 DIV in the presence of vehicle control (DMSO) or increasing concentrations of EZH1/2 inhibitor UNC1999. H3 was used as a loading control. **C** Heatmap depicting differentially expressed genes in response to EZH1/2 inhibition after treatment with 100 nM UNC1999, compared to DMSO. **D** The percent of genes in each PTM category (based on P7 GCP ChIPseq) that were up- or downregulated in cultured GCs in response to UNC1999 treatment, compared to DMSO. More than 10% of all bivalent genes were differentially expressed and the differentially expressed bivalent genes were significantly more likely to be upregulated than downregulated. Statistical analysis indicates the difference between the distribution of up- and downregulated genes between each PTM group. Pairwise Fisher test was performed using the rstatix package and p-values were corrected for multiple comparisons using the fdr method. ** p < 0.01, *** p < 0.001, **** p < 0.0001. **E-G** Volcano plots depicting differentially expressed bivalent (**E**), H3K4me3-only (**F**), or H3K27me3-only (**G**) genes in response to UNC1999 treatment, compared to DMSO. The number of up- or downregulated genes is indicated at the top of the chart. **H** Genome browser representation of RNAseq tracks for example bivalent genes at DIV5. *Actb* is shown as a control. **I** Volcano plot depicting differentially expressed genes (p-adjusted < 0.05, indicated with red) in response to UNC1999 treatment, compared to DMSO. Upregulated genes include several well-known marker genes of mature GCs, such as *Gabra6, Cbln3*, and *Grm4*. **J** Enriched GO categories of up- and downregulated genes after UNC1999 treatment compared to DMSO, identified using clusterProfiler package. The GO biological process categories were sorted by the adjusted P-value and the top 5 enriched non-redundant categories are shown for each group.

To investigate how the reduction in H3K27me3 impacts gene expression in developing GCs, we first used a well-established model of cultured GCs (Hatten, 1985). Cultured GC progenitors isolated from P6-P7 cerebella exit the cell cycle within 24 h (Tomoda et al., 1999), after which they differentiate, migrate, and begin maturation over 4-5 days *in vitro* (DIV) (Edmondson and Hatten, 1987; Solecki et al., 2004). To establish how gene expression in cultured GCs corresponds to *in vivo* GC development, we first used RNA-seq to determine the developmental gene expression pattern of cultured GCs at 0 DIV, 2 DIV, and 5 DIV. We identified the highest expressed genes at each time point (**Suppl. Fig. S4A-B**) and performed GO analysis to identify enriched biological processes. We found that categories enriched in the DIV0 transcriptome were associated with proliferation and DNA repair; categories enriched at DIV2 were associated with synaptic signaling and neuron projection morphogenesis; and categories enriched at DIV5 were associated with regulation of membrane potential (**Suppl. Fig. S4C**). These results were in line with the *in vivo* transcriptomics analysis (**Fig. 1C**) and suggested that the DIV0, DIV2, and DIV5 time points correspond to GC proliferation, differentiation, and maturation, respectively.

Next, we treated cultured GCs with UNC1999, an inhibitor of EZH1 and EZH2, to evaluate the outcome of EZH1/2 inhibition on GC development. H3K27me3 levels were reduced after treatment with increasing concentrations of UNC1999 for 5 DIV, compared with vehicle (DMSO) (**Fig. 6B**), and treatment with 100 nM UNC1999 for 5 DIV was used in subsequent experiments. In total, we identified 488 upregulated genes and 422 downregulated genes in response to UNC1999 treatment (**Fig. 6C**). While EZH1/2 inhibition was expected to reduce H3K27me3 levels at both bivalent and H3K27me3-only promoters, we found that the expression of bivalent genes was significantly more impacted by EZH1/2 inhibition than the expression of H3K4me3-only or H3K27me3-only genes (**Fig. 6D-G**). The majority of differentially expressed bivalent genes were upregulated after EZH1/2 inhibition (**Fig. 6D, H**), consistent with the primary effect of removing a repressive modification from these genes. Remarkably, some of the most significantly upregulated genes included well-known markers of mature GCs, including Gabra6, Neurod1, Cbln3, Grm4, and Grin2c, as well as the bivalent TFs Insm1, Neurod2, and Ebf3 identified above (**Fig. 6E**). Subsequent GO analysis of differentially expressed genes confirmed that downregulated genes included categories involved in cell growth and migration, whereas genes upregulated with UNC1999 treatment were involved in mature neuron function (**Fig. 6F**). Therefore, these results suggested that GCs cultured in the presence of an EZH1/2 inhibitor downregulate migration-related genes and upregulate genes involved in neuronal maturation.

To directly observe the effect of EZH1/2 inhibition on GC development, we treated *ex vivo* slices of P8 cerebellum with UNC1999 or vehicle control (DMSO) (**Fig. 7A**). Prior to UNC1999 or vehicle treatment, we electroporated P8 cerebella with a plasmid expressing a Venus fluorophore to allow visualization of the progression of labeled GCs through axon outgrowth, glial-guided migration and dendrite extension in specific layers of the cerebellum (Govek et al., 2018). In this system, after 60h *ex vivo*, GCs treated with the vehicle were located in the lower EGL, where they extended bipolar parallel processes, and in the molecular layer, where they exhibited a bipolar morphology with a single leading process typical of migrating cells and a trailing T-shaped axon (**Fig. 7B**, white arrows). In contrast, GCs treated with UNC1999 failed to enter the deeper aspects of the molecular layer (**Fig. 7B**). On average, the distance of migration, calculated by measuring the distance of the cell soma to the outer edge of the parallel fiber axons, was reduced by 25% after UNC1999 treatment, compared to the vehicle control (**Fig. 7C**). Instead, a significant number of UNC1999-treated cells displayed a multipolar morphology typical of post-migratory, terminally differentiated GCs (Zhu et al., 2016) (**Fig. 7B**, white asterisks), suggesting that inhibition of EZH1/2 induced premature maturation. Quantification of the number of bipolar and multipolar GCs in the molecular layer revealed that UNC1999 treatment significantly decreased the percentage of bipolar GCs and increased the percentage of multipolar GCs (**Fig. 7D**). Therefore, by both morphological criteria and position, UNC1999 treatment inhibited glial-guided migration of GCs and induced premature terminal differentiation. Together, these results suggest that disrupting bivalency in GCPs perturbs glial-guided neuronal migration, a key step in CNS development, and accelerates terminal differentiation.

**Figure 7.**
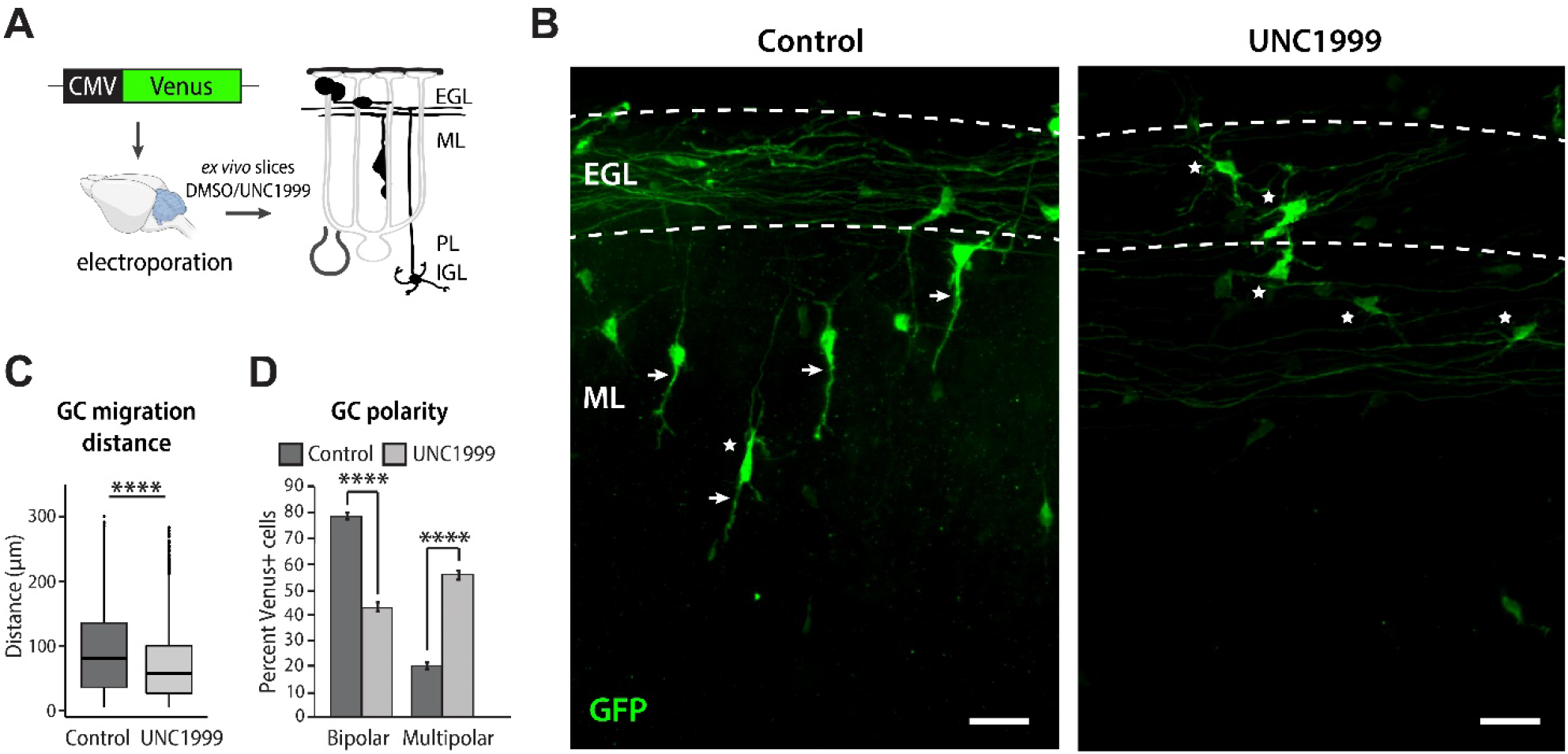
H3K4me3/H3K27me3 bivalency regulates the speed of GC neuronal maturation **A** Schematic of the experimental design. A plasmid encoding the Venus fluorophore was electroporated into the cerebellum and organotypic cerebellar slices were made and cultured in the presence of vehicle control (DMSO) or 200 nM UNC1999 for 60h. The morphology and subcortical localization of GCs was evaluated to identify GCs at different stages of development. **B** Images of electroporated GCs in *ex vivo* cerebellar slices, treated with vehicle control (DMSO) or UNC1999 for 60h. Control cells exhibit a bipolar morphology typical of migrating GCs, with a leading process oriented in the direction of migration (white arrows) and few multipolar cells (white asterisks). By contrast, cells treated with UNC1999 displayed inhibited migration and multipolar morphology. Scale bar, 25 μm. **C** GC migration distance after vehicle control (DMSO) or UNC1999 treatment, measured as the distance of the cell soma to the outer edge of the parallel fiber axons. Each replicate included slices from 2-3 postnatal P8 mouse pups per condition. GC migration distance was quantified from 4 replicates for a total of 868 cells (DMSO) or 2334 cells (UNC1999). **D** Quantification of the percentage of bipolar and multipolar GCs out of all Venus-expressing cells in the molecular layer. Each replicate included slices from 2-3 postnatal P8 mouse pups per condition. For the vehicle control (DMSO) condition, regions of interest were imaged from 5 replicates for a total of 593 cells. For the UNC1999 condition, regions of interest were imaged from 4 replicates for a total of 880 cells. Student’s t-test. **** p < 0.0001.

### GCP bivalent domains are enriched at marker genes of other cerebellar neurons

Given the proposed role of bivalency in lineage specification, we also analyzed whether bivalency is required to maintain cell-type-specific gene expression programs in proliferating GCPs and postmitotic neurons. To address this, we took advantage of published scRNA-seq data from adult mouse brain (Saunders et al., 2018), as well as the developmental lineage of cerebellar cells ((Vladoiu et al., 2019) **Suppl. Fig. S5A**). First, we reasoned that if bivalency is required to control lineage-appropriate gene expression in proliferating GCPs, we would observe a higher proportion of bivalent genes among genes specific to closely related cell types (e.g., unipolar brush cells), compared with distant cell types (e.g., microglia) (**Suppl. Fig. S5A**). To test this possibility, we evaluated the proportion of bivalent promoters identified in P7 GCPs among marker genes of different types of cerebellar cells, identified based on scRNA-seq (Saunders et al., 2018). Consistent with our hypothesis, a high proportion (35-50%) of marker genes of other cerebellar neurons were bivalent in proliferating GCPs, at a level that was significantly higher than expected genome-wide (**Suppl. Fig. S5A**). These results are consistent with the interpretation that bivalency maintains lineage-specific gene expression in committed neuronal progenitors.

To address the question of whether H3K27me3 and bivalency could repress the expression of other cell type marker genes in differentiated GCs, we analyzed how UNC1999 treatment impacts the expression of the identified marker genes in cultured GCs. We found that while GC-specific genes were significantly upregulated by UNC1999 treatment, the expression of other cerebellar cell type marker genes was essentially unchanged (**Suppl. Fig. S5B-D**). These results demonstrate that in postmitotic GCs, the loss of H3K27me3 does not lead to an overexpression of genes specific to other, even closely related, cell types. In other words, by the time immature neuronal progenitors become postmitotic, the loss of bivalency and H3K27me3 has minimal impact on the expression of genes specific to other cell lineages.

## DISCUSSION

Here we report that bivalency controls the gene expression programs required for glial-guided migration and maturation of an identified, well-studied CNS neuron, the cerebellar granule cell. Our studies using a genetically encoded probe for identifying bivalent domains demonstrate that bivalency exists in individual GCs. Our experiments using histone PTM and gene expression profiling demonstrate that bivalency regulates the expression of neuronal genes during GCP differentiation. In addition, inhibition of EZH1/2 inhibits glial-guided migration and accelerates dendrite formation, suggesting that bivalency regulates the timing of key steps in cerebellar GC differentiation. Therefore, these results demonstrate that the role of bivalency extends beyond the control of lineage development and pertains to the regulation of developmental progression of committed CNS progenitor cells.

Our study illustrates the importance of functional assays combined with molecular studies to define the relationship between gene expression and key biological processes. The use of a genetically encoded sensor with histone methylation reader domains for detecting bivalent nucleosomes is a powerful method for testing whether individual GCs have bivalent domains as opposed to bivalency being an averaged feature of GCs. Our data revealed discrete subnuclear focal areas of fluorescent labeling, consistent with the conclusion that individual GCs contain bivalent domains. These results were in line with a prior study utilizing the cMAP3 bivalency probe, which demonstrated that GCPs organize their poised chromatin in defined locations of H3.3-free euchromatin (Hoffman et al., 2020). The methodology does not provide quantitative details on the extent of bivalency in the positive domains, as discussed in (Delachat et al., 2018). However, a key finding from our experiments was that the fluorescent subnuclear foci were evident in proliferating GCPs and not in postmitotic GCs. This finding was consistent with the molecular data showing that bivalency peaked at P7, the time of maximal GCP proliferation.

Bivalent genes were primarily neuronal genes that were expressed at low to intermediate levels and were differentially expressed during GC differentiation and maturation. Of the genes identified as bivalent in the current study, of particular interest were several TFs known to be essential for specifying GC identity and differentiation. The finding that these genes also had the widest H3K4me3 peaks is consistent with prior studies showing that broad H3K4me3 domains mark genes that determine cell identity (Benayoun et al., 2014). Taken together, these findings confirmed the presence of bivalency in individual GCPs and implicated bivalency in the establishment of GC identity, the fine-tuning of gene expression levels, and the control of gene expression dynamics during neuronal development.

The role of chromatin dynamics in regulating gene expression demonstrated in the current study is consistent with our prior finding that the expression of *Tet* genes controls the expression of axon guidance and ion channel genes, underlying the formation of dendrites and synaptic connections in GCs (Zhu et al., 2016). TET enzymes catalyze the conversion of DNA 5-methylcytosine to 5-hydroxymethylcytosine, a modification that during terminal differentiation accumulates in neuronal chromatin in a cell-type specific manner and correlates with active transcription (Gabel et al., 2015; Kriaucionis and Heintz, 2009; Mellen et al., 2017; Stoyanova et al., 2021). In addition to regulating gene expression during GC maturation, TET activity is also required for the terminal differentiation of Purkinje cells (Stoyanova et al., 2021) and activity-dependent gene expression in hippocampal and cortical neurons (Rudenko et al., 2013). It is noteworthy that in mouse ESCs, TET1 directly interacts with PRC2 (Neri et al., 2013) and binds bivalent promoters (Villasenor et al., 2020), where it regulates the levels of DNA methylation (Gu et al., 2018; Verma et al., 2018; Xiang et al., 2020). Thus, a possible model emerges that in immature neurons, bivalency could mark the promoters that become targets of TET enzymes during neuronal differentiation, thereby orchestrating the formation of the regulatory chromatin landscape in mature neurons. The detailed characterization of the interactions between bivalency and DNA hydroxymethylation during neuronal development and circuitry formation will be important to resolve this question.

The importance of bivalency on GCP gene expression and development was revealed by treating *ex vivo* slices of cerebellar cortex and cultured GCs with UNC1999, a selective inhibitor for the H3K27 methyltransferases EZH1 and EZH2. Strikingly, UNC1999 inhibited migration and promoted dendrite extension of GCs in the organotypic slice assay. This result was supported by molecular studies in cultured GCs showing that GO pathways for migration were downregulated, whereas those for synapse formation were upregulated after treatment with UNC1999. Although inhibition of EZH1 and EZH2 was expected to alter both bivalent and H3K27me3 monovalent genes, we found that UNC1999 predominantly affected the expression of bivalent genes. These results were consistent with our observations that the levels of H3K27me3 are lower at bivalent genes than at H3K27me3-only genes, and that the ratio of H3K4me3 to H3K27me3 at bivalent domains correlates with gene expression levels. Further studies specifically targeting EZH1 and EZH2, as well as changes in the composition of the PRC2 subunits will be required to define how these two methyltransferases contribute to the regulation of H3K27me3 and bivalency in developing neurons. Moreover, it remains to be determined how H3K4 methyltransferases, in particular MLL2/KMT2B (Hu et al., 2013), and histone demethylases regulate bivalency and the timing of GC developmental progression.

The finding that EZH1/2 inhibition accelerates maturation but inhibits glial-guided migration was unexpected, given that prior reports have showed that stimulating early steps in GCP differentiation, such as by inhibiting the transcription factor Zeb1 (Singh et al., 2016) or overexpressing NeuroD1 (Hanzel et al., 2019), induces polarization required for the onset of migration and promotes glial-guided migration. By contrast, inhibiting bivalency prematurely induced the gene expression program associated with mature neurons, as evident from the upregulation of synaptic genes and increased percentage of multipolar GCs after treatment with UNC1999. The accelerated maturation was accompanied with inhibited migration, as GCs treated with UNC1999 downregulated genes involved with migration and failed to migrate into the IGL. Our results are therefore consistent with the model that bivalency is required to poise the expression of genes involved in terminal differentiation until migration is completed, and premature induction of neuronal maturation blocks glial-guided migration.

While bivalency peaked at P7 during GCP proliferation, we also observed bivalent domains in mature GCs, leaving open the question of the role of bivalency in mature neurons. Although most bivalent genes resolved to monovalent during GC maturation, we identified over 1000 bivalent genes that either remained bivalent or became *de novo* bivalent in mature GCs. Our experiments using the EZH1/2 inhibitor do not support the hypothesis that bivalency is required to prevent transdifferentiation of postmitotic neurons into other types of cells. However, we found that genes that remained bivalent in mature GCs were involved in developmental processes and included growth factors, axon guidance genes, and transcription factors important for GCP specification in development (*Atoh1, Gli3, Mycn*). While the present study did not examine the function of bivalency in mature neurons that are integrated into the cerebellar circuitry, it will be of interest to determine whether bivalency at these genes allows their reactivation, as a response to neuronal damage, for example. Another point to raise in this context is that *Atoh1, Gli3* and *Mycn* are all upregulated in a subtype of medulloblastoma that originates from GCs. Therefore, dysregulation of bivalent genes in postmitotic neurons could potentially have relevance in cancer pathogenesis.

The model that emerges from this study is that bivalency controls the timing of the progression of steps that define the characteristics of specific CNS neurons. In the future, it will be important to examine the role of other histone PTMs in regulating the developmental progression of neuronal cell types of the CNS. Understanding how epigenetic mechanisms control the functional maturation of neurons will inform future studies on the formation of CNS circuitries and the emergence of neurodevelopmental disorders.

## METHODS

### Animals

All animal experiments were conducted in accordance with the United States Animal Welfare Act and the National Institutes of Health’s policy to ensure proper care and use of laboratory animals for research, and under established guidelines and supervision by the Institutional Animal Care and Use Committee (IACUC) of the Rockefeller University. Mice were housed in accredited facilities of the Association for Assessment of Laboratory Animal Care (AALAC) in accordance with the National Institutes of Health guidelines. All efforts were made to minimize the suffering of animals. Both male and female animals were included in the study. C57Bl/6J mice were purchased from the Jackson Laboratories. *Tg(NeuroD1-Egfp-L10a)* and *Tg(Atoh1-Egfp-L10a)* TRAP mice were a gift from Dr. Nathaniel Heintz. The mice were group-housed in a specific pathogen free stage with *ad libitum* access to food and water under a 12-hour light/dark cycle (ON/OFF 7AM/7PM). Bedding and nest material were changed weekly. Where appropriate, ARRIVE guidelines are followed for reporting animal research.

### Plasmids

The pCIG2-Venus plasmid was described in (Govek et al., 2018). A plasmid encoding the cMAP3 bivalency probe was a gift from David Solecki (Delachat et al., 2018; Hoffman et al., 2020). The cMAP3 sequence, consisting of Polycomb chromodomain (PCD), Emerald fluorophore and transcription factor TFIID subunit 3 plant homeodomain (TAF3), was subcloned into the pCIG2 vector. The pCIG2-cMAP3 plasmid was further modified to co-express the tdTomato fluorophore under an IRES.

### Nuclei isolation and labeling

Nuclei isolation was adapted from (Xu et al., 2018) with modifications. P12 and P21 cerebella were dissected from C57Bl/6J mice and either snap-frozen or processed fresh. For nuclei isolation, 4-6 cerebella were pooled. The tissue was homogenized in homogenization buffer (0.25 M sucrose, 150 mM KCl, 5 mM MgCl2, 20 mM Tricine-KOH pH 7.8, 0.15 mM spermine, 0.5 mM spermidine, EDTA-free protease inhibitor cocktail, PhosSTOP, 1 mM DTT) by 30 strokes with loose (A) pestle followed by 30 strokes with tight (B) pestle in a glass Dounce homogenizer on ice. The homogenate was supplemented with 0.92 volume of iodixanol solution (50% Iodixanol/Optiprep, 150 mM KCl, 5 mM MgCl2, 20 mM Tricine pH 7.8, 0.15 mM spermine, 0.5 mM spermidine, EDTA-free protease inhibitor cocktail, PhosSTOP, 1 mM DTT) and laid on a 27% iodixanol cushion. Nuclei were pelleted by centrifugation for 30 min at 17,200*g* at 4 °C in a standard tabletop centrifuge. The pellet was resuspended in the homogenization buffer and passed through a cell strainer. Resuspended nuclei were fixed with 1% formaldehyde for 7 min at room temperature and quenched with 0.125M glycine for 5 min at room temperature. Nuclei were pelleted by centrifugation for 5 min at 1000*g* at 4 °C and washed once with the homogenization buffer and once with wash buffer (3% BSA, 0.05% Triton X-100 in PBS). Nuclei were blocked with block buffer (6% BSA, 0.05% Triton X-100 in PBS) for 1h at room temperature and incubated with anti-NeuN (Millipore #MAB377, RRID:AB_2298772, 1:500-1:1000) and anti-NeuroD (Santa Cruz Biotechnology #sc-1084, RRID:AB_630922, 1:100) antibodies at 4 °C o/n. The nuclei were washed three times with wash buffer for 5 min at 1000*g* at 4 °C centrifugation in between, incubated with secondary antibodies (Invitrogen Alexa Fluor 488 Donkey anti-Mouse IgG #A21202, RRID: AB_141607 and Invitrogen Alexa Fluor 555 Donkey anti-Rabbit IgG #A31572, RRID: AB_162543, 1:500) for 30 min at room temperature, and washed three times with wash buffer. Primary and secondary antibodies were diluted in the block buffer.

### Flow cytometry

Labeled nuclei were resuspended in 1% BSA, 10 mM HEPES in PBS, and incubated with DyeCycle Ruby (Thermo Fisher Scientific #V10309) for 30 min at room temperature protected from light. Sorting was carried out at the Flow Cytometry Resource Center at Rockefeller University, using a BD FACSAria cell sorter using the 70 μm nozzle and 488 nm, 561 nm, and 635/640 nm lasers. Samples were gated using DyeCycle Ruby to identify singlets, followed by the sorting of NeuN+/NeuroD1+ population. Sorted nuclei were pelleted by centrifugation, resuspended in DPBS to divide into aliquots of 10 × 10^6^ nuclei/tube, centrifuged again, and stored at −80 °C. In these experiments, 95-98% of singlets were NeuN+/NeuroD1+ and the average yield was 6-10 × 10^6^ nuclei per one P12 or P21 cerebellum. The specificity of the primary antibodies and the gating strategy was confirmed using *Tg(NeuroD1-Egfp-L10a)* mice that express EGFP specifically in postmitotic GCs of the cerebellum.

### GCP purification

P7 GCPs were isolated from C57Bl/6J mice as previously described (Hatten, 1985). Briefly, isolated cerebella were dissected in ice-cold CMF-PBS and dissociated with trypsin-DNase I for 5 min at 37 °C, followed by centrifugation for 5 min at 700*g* at 4 °C. Trypsin-DNase I was removed, and the cells were triturated in DNase-CMF-PBS 10x with a transfer pipette, 10x with a fine fire-polished Pasteur pipette, and 10x with an extra-fine Pasteur pipette. The cell homogenate was then centrifuged for 5 min at 700*g* at 4 °C and the cell pellet was resuspended in DNase-CMF-PBS. The cells were subjected to Percoll gradient sedimentation to enrich for proliferating GCPs, and subsequently pre-plated for 15-30 min on a Petri dish and 1-2h on a tissue culture dish to remove glia and fibroblasts. The medium was collected and centrifuged for 5 min at 700*g* at 4 °C to pellet the GCPs. Cells dedicated for ChIP were subsequently resuspended in DPBS, fixed with 1% formaldehyde for 7 min at room temperature, quenched with 0.125M glycine for 5 min, washed once with DPBS, and frozen in aliquots of 5-10 × 10^6^ cells.

#### GCP cultures and UNC1999 treatment

GCPs isolated after pre-plating were subsequently plated on a 24-well plate precoated with poly-D-lysine (0.1 mg/ml, Sigma #P1024) and Matrigel in granule cell medium (BME (Life Technologies # 21010-046), 2mM L-Glutamine (Gibco #25030-016), 1X Pen-Strep (Gibco #15140-015), 0.9% glucose (Sigma #G8769), 10% horse serum (Gibco #16050-122, heat-inactivated) and 5% fetal bovine serum (Gibco #26140-079, heat-inactivated)) at a final concentration of 2.5 × 10^6^ cells in 600 μl. UNC1999 (Tocris Biosciences, final concentration 10-100 nM) or an equivalent volume of DMSO was added to the culture medium starting at 0 DIV. One half of culture medium was replaced with fresh medium plus DMSO or UNC1999 daily until 5 DIV. Cells were collected by scraping, pelleted by centrifugation, and stored at −80°C until processed for RNA and protein isolation.

### RNA isolation

RNA from cultured GCs was isolated using RNeasy Micro Kit (Qiagen #74004) with on-column DNA digestion. RNA quality was analyzed with the Bioanalyzer RNA 6000 Pico kit (Agilent) prior to library preparation.

### Protein isolation

Nuclear and cytoplasmic protein fractions were isolated using subcellular fractionation protocol. Briefly, nuclei were isolated in subcellular fractionation buffer (20 mM HEPES, 10 mM KCl, 2 mM MgCl2, 1 mM EDTA, 1 mM EGTA, supplemented with 0.5 mM DTT, 0.2 mM PMSF, and protease and phosphatase inhibitors), incubated on ice for 15 min, passed through a 27G needle 3-5 times, incubated on ice for another 20 min and pelleted for 5 min at 3000 rpm at 4°C. Supernatant containing the cytoplasmic fraction was stored at −80°C. The pellet containing the nuclei was then resuspended in TBS containing 0.1% SDS and sonicated using Bioruptor Pico (Diagenode) on at 30 seconds on–30 seconds off for 10 cycles. Insoluble fraction was pelleted by centrifugation for 10 min at 13000 rpm at 4°C and supernatant containing nuclear proteins was stored at −80°C.

### Immunoblotting

Samples were run on 16% Tris-glycine SDS-PAGE gels (Invitrogen) and transferred to 0.2 μm nitrocellulose membranes. The membranes were blocked in 5% milk in TBST and incubated with antibody solutions in 2.5% milk, with TBST washes in-between. Proteins were visualized using Immobilon ECL (Invitrogen) and D1001 KwikQuant Imager (Kindle Biosciences). The following primary antibodies were used: Rabbit anti-H3K27me3 (Cell Signaling Technology #9733, RRID:AB_2616029, 1:300) and Rabbit anti-H3 (Abcam #ab1791, RRID:AB_302613, 1:5000).

### Chromatin immunoprecipitation (ChIP)

#### Lysis and sonication

Fixed GCPs were thawed on ice, resuspended in Cell Lysis Buffer (50 mM HEPES-KOH pH 7.5, 140 mM NaCl, 1 mM EDTA, 10% glycerol, 0.5% NP-40, and 0.25% Triton X-100, supplemented with 0.5 mM DTT, 0.2 mM PMSF, and protease and phosphatase inhibitors; 1 ml per 10 × 10^6^ cells) and incubated for 10 minutes at 4 °C using end-over-end rotation. Nuclei were isolated by centrifugation at 1350*g* for 5 minutes at 4 °C. Fixed and sorted nuclei from P12 and P21 mice were thawed on ice. GCP nuclei and nuclei isolated with FACS were subsequently resuspended in Nuclei Lysis Buffer (50 mM Tris-HCl pH 8, 10 mM EDTA, and 1% SDS, supplemented with 0.5 mM DTT, 0.2 mM PMSF, and protease and phosphatase inhibitors; 140 μl per 10 × 10^6^ nuclei) and incubated for 10 minutes at RT using end-over-end rotation. Lysates were passed through a 27G needle 2-3 times and sonicated using a Covaris E220 focused ultrasonicator in an AFA Fiber Pre-Slit Snap-Cap microTUBE at 140W peak power, 10% duty factor, and 200 cycles/burst for 1 min 50 s (P7 GCP nuclei) or 2 min 10 s (P12 and P21 GC nuclei). Triton X-100 was added at a final concentration of 1% and insoluble chromatin was pelleted by centrifugation at 13,000 rpm for 10 minutes at 4 °C. 5% of chromatin was used for determining sonication efficiency. Samples were stored at −80 °C until processing.

#### CHIP

Sonicated chromatin was diluted 10x with Dilution Buffer (1 mM Tris-HCl pH 8.0, 167 mM NaCl, 0.1 mM EDTA, 0.5% sodium deoxycholate, and 1% NP-40, supplemented with 0.5 mM DTT, 0.2 mM PMSF, and protease and phosphatase inhibitors). In a subset of experiments, Drosophila chromatin (Active Motif #53083; 1 ng per 1 μg chromatin) was added to the samples. 5 % of chromatin was used for input. Protein A Dynabeads (Invitrogen #10002D) were coated with antibodies (Cell Signaling Technology Rabbit anti-H3K27me3 Cat#9733, RRID:AB_2616029 and Active Motif Rabbit anti-H3K4me3 Cat# 39159, RRID:AB_2615077) in 0.5% BSA in PBS for 2 hours at 4 °C using end-to-end rotation and washed 3 times with 0.5% BSA in PBS. Antibody-coated beads were added to the samples and incubated at 4 °C rotating overnight. The beads were then washed 8 times with Wash Buffer (50 mM HEPES-KOH pH 7.6, 500 mM LiCl, 1 mM EDTA, 1% NP-40, and 0.7% sodium deoxycholate) and once with TE containing 50 mM NaCl. The chromatin was eluted from the beads with Elution Buffer (50 mM Tris-HCl pH 8.0, 10 mM EDTA, and 1% SDS) for 30 minutes shaking at 65 °C and samples were incubated at 65 °C overnight to reverse crosslinks. RNA was digested with DNase-free RNase (5 μg/ml, Roche #11119915001) for 1 hour at 37 °C, and proteins were digested with proteinase K (0.2 mg/ml, Thermo Scientific #EO0491) for 1 hour at 55 °C. DNA was purified using PCR Purification Kit (Qiagen #28104). ChIP efficiency prior to sequencing was evaluated using ChIP qPCR (not shown).

### ChIP sequencing

ChIP-seq libraries were prepared using the TruSeq ChIP Library Preparation Kit (Illumina #IP-202-1012 or #IP-202-1024) or the NEBNext Ultra II DNA Library Prep Kit for Illumina (E7645S). The quality of the sequencing libraries was evaluated using the Agilent 2200 TapeStation with D1000 High Sensitivity ScreenTape. The samples were sequenced at the Rockefeller University Genomics Resource Center using a NextSeq 500 sequencer (Illumina) to obtain 75 bp single-end reads.

### Translating ribosome affinity purification (TRAP)

#### Antibody binding

TRAP was performed as previously described (Doyle et al., 2008; Heiman et al., 2008). For each reaction, 300 μl of Streptavidin MyOne T1 Dynabeads (Invitrogen #65601) were washed 5x with 1x PBS and incubated with 120 μg biotinylated protein L (Pierce #29997, reconstituted at 1 μg/μl in PBS) in 1x PBS in a total volume of 1 ml for 35 minutes at RT, using end-over-end rotation. The beads were then washed 5 times with 3% IgG, Protease-free BSA (JacksonImmuno #001-000-162) in 1x PBS and subsequently incubated with 50 μg each of 19C8 and 19F7 anti-GFP monoclonal antibodies (Memorial Sloan-Kettering Monoclonal Antibody Facility) in 500 μl of 0.15 M KCl TRAP Wash Buffer (10 mM HEPES-KOH pH 7.4, 5 mM MgCl2, 150 mM KCl, and 1% NP-40, supplemented with 100 μg/ml cycloheximide (Millipore Sigma #C7698-1g) in methanol, 0.5 mM DTT (Thermo Fisher Scientific #R0861), and 20 U/ml RNasin (Fisher Scientific #PR-N2515) just before use), for 30 min using end-over-end rotation. After antibody binding, the beads were washed 3 times with 0.15 M KCl TRAP Wash Buffer, resuspended in 0.15 M KCl TRAP Wash Buffer, and each reaction was supplemented with 30 mM DHPC (Avanti #850306P).

#### Brain lysate preparation

Cerebella from P7, P12, or P21 *Tg(NeuroD1-Egfp-L10a)* mice were dissected and placed in TRAP Dissection Buffer (2.5 mM HEPES-KOH pH 7.4, 35 mM glucose and 4 mM NaHCO3 in 1x HBSS, supplemented with 100 μg/ml cycloheximide) until ready for homogenization. The tissue was homogenized in chilled TRAP Lysis Buffer (10 mM HEPES-KOH pH 7.4, 5 mM MgCl2, and 150 mM KCl, supplemented with 0.5 mM DTT, 100 μg/ml cycloheximide, protease inhibitor cocktail (Sigma #11836170001), 40 U/ml RNasin and 20 U/ml Superasin (Thermo Fisher Scientific #AM2694) just before use) using a motor-driven Teflon-Glass homogenizer at 900 rpm, 12 strokes. The homogenate was centrifuged at 2,000*g* for 10 minutes at 4 °C and 3% of the supernatant was set aside as an input. The rest of the supernatant was mixed with NP-40 to a final concentration of 1% and DHPC to a final concentration of 30 mM and incubated on ice for 5 minutes. Subsequently, the samples were centrifuged at 20,000*g* for 10 minutes at 4 °C and the supernatant was used for immunoprecipitation.

#### GCPpreparation for TRAP

GCPs from P7 *Tg(Atoh1-Egfp-L10a)* mice were isolated using Percoll gradient centrifugation as described above. Pooled GCPs from 2 to 3 mice were homogenized in TRAP Lysis Buffer and processed in parallel to brain lysates described above. Since the isolated cells were enriched for GCPs, input was not collected.

#### Immunoprecipitation

The samples were incubated with GFP-conjugated beads o/n at 4 °C with end-to-end rotation. Subsequently, the beads were washed 4 times in 0.35 mM KCl TRAP Wash Buffer (10 mM HEPES-KOH pH 7.4, 5 mM MgCl2, 350 mM KCl, and 1% NP-40, supplemented with 100 μg/ml cycloheximide, 0.5 mM DTT, and 20 U/ml RNasin just before use) and resuspended in 100 μl RLT Buffer from the RNeasy Micro Kit (Qiagen #74004) supplemented with 1% β-mercaptoethanol. The resuspended beads were incubated at RT for 10 min, placed on a magnet, and the supernatant containing the RNA was collected and purified using the RNeasy Micro Kit. RNA integrity was determined using Bioanalyzer RNA 6000 Pico kit (Agilent) prior to library preparation.

### RNA sequencing

RNA-seq libraries were prepared using the NEBNext Ultra II RNA Library Prep Kit for Illumina (NEB #E7770S) in conjunction with the NEBNext Poly(A) mRNA Magnetic Isolation Module (NEB #E7490) and NEBNext Multiplex Oligos for Illumina (NEB #E7335, #E7500). 10 ng (TRAP RNA from *Tg(Atoh1-Egfp-L10a)* mice), 800 ng (TRAP RNA from *Tg(NeuroD1-Egfp-L10a)* mice) or 500 ng (total RNA from cultured GCs) of RNA was used for library preparation. Input samples for each developmental age in the TRAP experiment were pooled in equal amounts to yield one input per time point. The quality of the sequencing libraries was evaluated using the Agilent 2200 TapeStation with D1000 High Sensitivity ScreenTape. The samples were sequenced at the Rockefeller University Genomics Resource Center using a NextSeq 500 sequencer (Illumina) to obtain 75 bp single-end reads.

### Bioinformatics analysis

Sequence and transcript coordinates for mouse mm10 UCSC genome and gene models were retrieved from the Bioconductor Bsgenome.Mmusculus.UCSC.mm10 (version 1.4.0) and TxDb.Mmusculus.UCSC.mm10.knownGene (version 3.4.0) libraries, respectively.

ChIP-seq reads were mapped using the Rsubread package’s align function (version 1.30.6) (Liao et al., 2019). Peak calls were made with HOMER (version 4.11, style histone, size=1000, minDist=2500) (Heinz et al., 2010). Consensus peaks were determined to be peaks that were found in the majority of replicates. Peaks were annotated and genome distribution was determined using the ChIPseeker package (version 1.30.3) (Yu et al., 2015). Bivalent genes were those assessed to contain overlapping H3K27me3 and H3K4me3 peaks that overlap with the TSS (+/- 500bp). We note that the peaks of H3K27me3 and H3K4me3 signals at identified bivalent regions did not always coincide. Pairwise comparisons between developmental timepoints were made using DEseq2 (version 1.34.0) (Love et al., 2016) using counts from TSS with significant genes considered as (padj < 0.05). DESeq2 was also used to derive the bivalency ratio, through a pairwise comparison between H3K27me3 and H3K4me3 at TSS. Normalized, fragment extended signal bigWigs were created using the rtracklayer package (version 1.40.6) (Lawrence et al., 2009), and then visualized and exported using Gviz (version 1.38.4). Heatmaps and metaplots were generated with profileplyr (version 1.10.2) (Carroll and Barrows, 2022).

RNA-seq reads were mapped using Rsubread. Transcript expressions were calculated using the Salmon quantification software (version 0.8.2) (Patro et al., 2017) and gene expression levels as TPMs and counts were retrieved using Tximport (version 1.8.0) (Love et al., 2016). Normalization and rlog transformation of raw read counts in genes, PCA and differential gene expression analysis were performed using DESeq2. Additional sample to sample variability assessment was made with heatmaps of between-sample distances using pheatmap (version 1.0.12). Pairwise comparisons between developmental timepoints were made using DEseq2 with significant genes considered as (padj < 0.01, log_2_FC≥1). For the TRAP dataset, highly depleted genes (TRAP/input < 0.6 in at least three developmental time points) were excluded prior to enrichment analysis. For both ChIP-seq and RNA-seq, GO enrichment analysis was performed using clusterProfiler (version 4.2.2) (Yu et al., 2012). General processing and exploration of data were performed using the tidyverse packages (Wickham et al., 2019). Plots were prepared using ggplot2 (version 3.3.6) (Wickham, 2016), ggVennDiagram (version 1.2.2) (Gao et al., 2021) and ggalluvial (version 0.12.3) (Brunson, 2020).

For the identification of cerebellar cell type specific genes, we used data from a previously published scRNA-seq dataset from the adult mouse brain (Saunders et al., 2018). Subclusters were combined as necessary to obtain one metacell per each major cerebellar cell type (granule cells, Purkinje cells, unipolar brush cells, stellate/basket cells, Golgi cells, astrocytes/Bergmann glia, oligodendrocytes, microglia, endothelial cells and pericytes/mural cells). To obtain a list of marker genes for each metacell, normalized counts for each cell type were scaled by z-score, sorted, and cut-off values were identified using the elbow point method. For each cell type, only genes that had a z-score of at least 1.5x higher than the next highest were included. The identified cell type marker genes are indicated in Supplementary Table S2.

### *Ex vivo* organotypic slices

P8 cerebella were dissected from C57BL/6J mouse brains in HBSS containing 2.5 mM HEPES (pH 7.4), 46 mM D-glucose, 1 mM CaCl2, 1 mM MgSO_4_ and 4 mM NaHCO3 (hereto referred to as HBSS with extra glucose) on ice. The dissection medium was then removed and pCIG-Venus or pCIG-cMAP3-tdTomato DNA was diluted to 0.5 μg/μl in HBSS with extra glucose. The cerebella were soaked in the DNA for 15-20 minutes on ice and were transferred one at a time into the well of an electroporation chamber (Protech International Inc. CUY520P5 platinum electrode L8xW5xH3 5mm gap) on ice. The cerebella were electroporated dorsal to ventral for 50 ms at 80 V, for a total of 5 pulses with an interval of 500 ms between pulses, using an electro-square-porator, ECM 830 (BTX Genetronics). The cerebella were then removed from the chamber, recovered in HBSS with extra glucose on ice for 10 minutes, embedded in 3% agarose in HBSS containing regular glucose (30 mM D-glucose), and 250 μm coronal slices were made using a Leica VT1000S vibratome set at a speed of 3 and frequency of 6. Slices were then placed on MILLICELL CM 0.4 μm culture plate inserts in a 6-well plate with 1.5 ml of culture medium (BME, 25 mM D-glucose, 1x Glutamine, 1x ITS and 1x Pen-Strep) below the insert. For the cMAP3 experiments, slices were incubated at 35°C/5% CO2 for 6, 12, 48 and 72h before fixing with 4% PFA/4% sucrose/PBS for 2h at room temperature. For the UNC1999 experiments, 200 nM UNC1999 in DMSO (or an equivalent volume of DMSO for the control conditions) was added to the culture medium just prior to plating the cerebellum. The organotypic slices were incubated at 35°C/5% CO2 for 60 hours before fixing with 4% PFA/4% sucrose/PBS for 2h at room temperature.

### Immunostaining and imaging

#### Immunostaining

For immunostaining of organotypic slices, fixed slices were washed three times in PBS for 30 minutes each time at room temperature, and were permeabilized and blocked overnight in 1x PBS/0.3% Triton X-100/10% Normal Donkey Serum (NDS) at 4°C. The sections for cMAP3 experiments were then incubated overnight at 4°C with rabbit anti-GFP (Thermo Fisher Scientific Cat# A-11122, RRID:AB_221569, 1:2000), goat anti-tdTomato (SICGEN Cat# AB8181-200, 1:200) and mouse anti-Calbindin (CALB1) (Swant Cat# 300, RRID:AB_10000347, 1:1000) primary antibodies diluted in 1x PBS/0.3% Triton X-100/10% NDS. The sections for the UNC1999 experiments were then incubated overnight at 4°C with rabbit anti-GFP (Thermo Fisher Scientific Cat# A-11122, RRID:AB_221569, 1:2000) and mouse anti-Calbindin (CALB1) (Swant Cat# 300, RRID:AB_10000347, 1:1000) primary antibodies diluted in 1x PBS/0.3% Triton X-100/10% NDS. Subsequently, the sections were washed 3 times for 20 minutes each time in 1x PBS/0.3% Triton X-100/10% NDS at room temperature and incubated overnight at 4°C with secondary antibodies (Thermo Fisher Scientific Cat# A-21206, RRID:AB_2535792, 1:500; Thermo Fisher Scientific Cat# A-21203, RRID:AB_141633, 1:500; Thermo Fisher Scientific Cat# A-11058, RRID:AB_2534105, 1:500; Abcam Cat# ab175658, RRID:AB_2687445, 1:500) diluted in 1x PBS/0.3% Triton X-100/10% NDS. The sections were then washed 4 times for 30 minutes each time, adding DAPI to the next to last wash of the slices for the UNC1999 experiments, the remaining agarose was carefully removed, and the sections were mounted with Molecular Probes ProLong Gold anti-fade mounting media (Invitrogen #P36934).

#### Imaging

The slices were imaged with a Carl Zeiss Axiovert 200M/Perkin Elmer Ultraview spinning disk confocal microscope equipped with a 25x objective, and images were acquired and viewed using Velocity software.

#### Quantification

Images were exported from Velocity software and figures were made in Adobe Photoshop and Illustrator. For the UNC1999 experiments, cell number was counted manually using the “count” tool and the distance of migration was measured manually using the “ruler” tool in Photoshop. Venus-positive cells with two processes emanating from the cell soma were considered bipolar and those with more than two processes emanating from the soma were considered multipolar. Co-immunostaining and imaging the slices with mouse anti-Calbindin to visualize the Purkinje cell layer (PCL) and DAPI to define the EGL and IGL aided in defining the EGL, ML, and PCL of the cerebellar cortex. The percentage of bipolar and multipolar GCs in the ML was calculated out of the total number of Venus-positive cells (n=593 cells for the control condition and n=880 cells for the UNC1999 condition). Cells for which polarity could not be determined were excluded from the analysis. The distance of migration was determined by measuring the distance from the cell soma to the outermost edge of the parallel fiber track for regions of interest in lobes on the surface of the slice. For regions of interest in internal areas of a slice, where two lobes and their parallel fiber tracks were adjacent, the outermost edge of each parallel fiber track was estimated to be located ½ the distance between the Purkinje cell layers for each lobe. Each replicate included slices from 2-3 postnatal P8 mouse pups per condition. For the control condition, regions of interest were imaged from 5 replicates and for the UNC1999 condition, regions of interest were imaged from 4 replicates.

### Quantification and statistical analysis

Sample size was determined based on prior studies. No statistical methods were used to predetermine sample size. Data was excluded based on insufficient quality (assessed immediately after acquisition and before analysis). Statistical tests to determine outliers were not performed and outliers were not excluded from the analysis. Data on bar graphs is presented as mean ± standard error of the mean. Data on box plots indicates median, first and third quartiles (lower and upper hinges) and smallest and largest observations (lower and upper whiskers, excluding outliers). Statistical analyses were performed in R, using the stats and rstatix packages. Pairwise comparisons were made using unpaired Student’s t-test. Multiple comparisons were made with analysis of variance (ANOVA), followed by Tukey HSD *post hoc* test, or, where appropriate, with pairwise t-test followed by correction for multiple comparisons with the Benjamini&Hochberg (BH)/fdr method. Non-parametric data were analyzed with the Kruskal-Wallis test, followed by Dunn’s *post hoc* test. Comparisons between proportions were performed using Fisher’s exact test, correcting for multiple comparisons using the fdr method. The level of statistical significance was set at P < 0.05.

## Data availability

ChIP-seq and RNA-seq datasets are deposited to GEO under accession number GSE223487.

## Acknowledgments

We are grateful to the members of the Hatten and Allis laboratories, Dr. Nathaniel Heintz, and Dr. Erica Korb for helpful discussions throughout the project. We thank Yin Fang, Katherine Dinan and Yana Leshchynska for technical assistance and Dr. David J. Solecki for the kind gift of the cMAP3 bivalency probe. We also thank Dr. Ningyan Cheng, Katherine Dinan, Danielle Keahi, Dr. Kert Mätlik, and Dr. David Solecki for critical comments on the manuscript. We gratefully acknowledge the Bioinformatics Resource Center, especially the advice of Dr. Thomas Carroll, Flow Cytometry Resource Center, Genomics Resource Center, and the Comparative Bioscience Center at the Rockefeller University for advice and support. Schematics were prepared with the help of BioRender. K.M. was supported by fellowships from the Sigrid Juselius Foundation and the Leon Levy Foundation. This work was also supported by NIH grant R21 NS114545 (M.E.H.).

## Author contributions

Conceptualization: K.M., M.E.H., Formal Analysis: K.M., M.R.P., E.E.G., Investigation: K.M., E.E.G., Resources: M.E.H., C.D.A., Visualization: K.M., E.E.G., Writing - original draft: K.M., Writing - review and editing: E.E.G., M.R.P., M.E.H., Supervision: M.E.H., Funding acquisition: K.M., M.E.H.

## References

Aoyama, K., Oshima, M., Koide, S., Suzuki, E., Mochizuki-Kashio, M., Kato, Y., Tara, S., Shinoda, D., Hiura, N., Nakajima-Takagi, Y., et al. (2018). Ezh1 Targets Bivalent Genes to Maintain Self-Renewing Stem Cells in Ezh2-Insufficient Myelodysplastic Syndrome. iScience 9, 161–174.

Barski, A., Cuddapah, S., Cui, K., Roh, T.Y., Schones, D.E., Wang, Z., Wei, G., Chepelev, I., and Zhao, K. (2007). High-resolution profiling of histone methylations in the human genome. Cell 129, 823–837.

Behesti, H., Kocabas, A., Buchholz, D.E., Carroll, T.S., and Hatten, M.E. (2021). Altered temporal sequence of transcriptional regulators in the generation of human cerebellar granule cells. Elife 10.

Benayoun, B.A., Pollina, E.A., Ucar, D., Mahmoudi, S., Karra, K., Wong, E.D., Devarajan, K., Daugherty, A.C., Kundaje, A.B., Mancini, E., et al. (2014). H3K4me3 breadth is linked to cell identity and transcriptional consistency. Cell 158, 673–688.

Bernstein, B.E., Kamal, M., Lindblad-Toh, K., Bekiranov, S., Bailey, D.K., Huebert, D.J., McMahon, S., Karlsson, E.K., Kulbokas, E.J., 3rd, Gingeras, T.R., et al. (2005). Genomic maps and comparative analysis of histone modifications in human and mouse. Cell 120, 169–181.

Bernstein, B.E., Mikkelsen, T.S., Xie, X., Kamal, M., Huebert, D.J., Cuff, J., Fry, B., Meissner, A., Wernig, M., Plath, K., et al. (2006). A bivalent chromatin structure marks key developmental genes in embryonic stem cells. Cell 125, 315–326.

Brickley, S.G., Cull-Candy, S.G., and Farrant, M. (1996). Development of a tonic form of synaptic inhibition in rat cerebellar granule cells resulting from persistent activation of GABAA receptors. J Physiol 497 (Pt 3), 753–759.

Brunson, J.C. (2020). ggalluvial: Layered Grammar for Alluvial Plots. Journal of Open Source Software 5, 2017.

Carroll, T., and Barrows, D. (2022). profileplyr: Visualization and annotation of read signal over genomic ranges with profileplyr.

Cenik, B.K., and Shilatifard, A. (2021). COMPASS and SWI/SNF complexes in development and disease. Nat Rev Genet 22, 38–58.

Consalez, G.G., Goldowitz, D., Casoni, F., and Hawkes, R. (2020). Origins, Development, and Compartmentation of the Granule Cells of the Cerebellum. Front Neural Circuits 14, 611841.

Delachat, A.M., Guidotti, N., Bachmann, A.L., Meireles-Filho, A.C.A., Pick, H., Lechner, C.C., Deluz, C., Deplancke, B., Suter, D.M., and Fierz, B. (2018). Engineered Multivalent Sensors to Detect Coexisting Histone Modifications in Living Stem Cells. Cell Chem Biol 25, 51–56 e56.

Dixon, J.R., Jung, I., Selvaraj, S., Shen, Y., Antosiewicz-Bourget, J.E., Lee, A.Y., Ye, Z., Kim, A., Rajagopal, N., Xie, W., et al. (2015). Chromatin architecture reorganization during stem cell differentiation. Nature 518, 331–336.

Doyle, J.P., Dougherty, J.D., Heiman, M., Schmidt, E.F., Stevens, T.R., Ma, G., Bupp, S., Shrestha, P., Shah, R.D., Doughty, M.L., et al. (2008). Application of a translational profiling approach for the comparative analysis of CNS cell types. Cell 135, 749–762.

Edmondson, J.C., and Hatten, M.E. (1987). Glial-guided granule neuron migration in vitro: a high-resolution time-lapse video microscopic study. J Neurosci 7, 1928–1934.

Espinosa, J.S., and Luo, L. (2008). Timing neurogenesis and differentiation: insights from quantitative clonal analyses of cerebellar granule cells. J Neurosci 28, 2301–2312.

Feng, X., Juan, A.H., Wang, H.A., Ko, K.D., Zare, H., and Sartorelli, V. (2016). Polycomb Ezh2 controls the fate of GABAergic neurons in the embryonic cerebellum. Development 143, 1971–1980.

Gabel, H.W., Kinde, B., Stroud, H., Gilbert, C.S., Harmin, D.A., Kastan, N.R., Hemberg, M., Ebert, D.H., and Greenberg, M.E. (2015). Disruption of DNA-methylation-dependent long gene repression in Rett syndrome. Nature 522, 89–93.

Gao, C.H., Yu, G., and Cai, P. (2021). ggVennDiagram: An Intuitive, Easy-to-Use, and Highly Customizable R Package to Generate Venn Diagram. Front Genet 12, 706907.

Govek, E.E., Wu, Z., Acehan, D., Molina, H., Rivera, K., Zhu, X., Fang, Y., Tessier-Lavigne, M., and Hatten, M.E. (2018). Cdc42 Regulates Neuronal Polarity during Cerebellar Axon Formation and Glial-Guided Migration. iScience 1, 35–48.

Gu, T., Lin, X., Cullen, S.M., Luo, M., Jeong, M., Estecio, M., Shen, J., Hardikar, S., Sun, D., Su, J., et al. (2018). DNMT3A and TET1 cooperate to regulate promoter epigenetic landscapes in mouse embryonic stem cells. Genome Biol 19, 88.

Hanzel, M., Rook, V., and Wingate, R.J.T. (2019). Mitotic granule cell precursors undergo highly dynamic morphological transitions throughout the external germinal layer of the chick cerebellum. Sci Rep 9, 15218.

Hatten, M.E. (1985). Neuronal regulation of astroglial morphology and proliferation in vitro. J Cell Biol 100, 384–396.

Heiman, M., Schaefer, A., Gong, S., Peterson, J.D., Day, M., Ramsey, K.E., Suarez-Farinas, M., Schwarz, C., Stephan, D.A., Surmeier, D.J., et al. (2008). A translational profiling approach for the molecular characterization of CNS cell types. Cell 135, 738–748.

Heinz, S., Benner, C., Spann, N., Bertolino, E., Lin, Y.C., Laslo, P., Cheng, J.X., Murre, C., Singh, H., and Glass, C.K. (2010). Simple combinations of lineage-determining transcription factors prime cis-regulatory elements required for macrophage and B cell identities. Mol Cell 38, 576–589.

Hirabayashi, Y., Suzki, N., Tsuboi, M., Endo, T.A., Toyoda, T., Shinga, J., Koseki, H., Vidal, M., and Gotoh, Y. (2009). Polycomb limits the neurogenic competence of neural precursor cells to promote astrogenic fate transition. Neuron 63, 600–613.

Hoffman, D.P., Shtengel, G., Xu, C.S., Campbell, K.R., Freeman, M., Wang, L., Milkie, D.E., Pasolli, H.A., Iyer, N., Bogovic, J.A., et al. (2020). Correlative three-dimensional super-resolution and block-face electron microscopy of whole vitreously frozen cells. Science 367.

Horn, Z., Behesti, H., and Hatten, M.E. (2018). N-cadherin provides a cis and trans ligand for astrotactin that functions in glial-guided neuronal migration. Proc Natl Acad Sci U S A 115, 10556–10563.

Hu, D., Garruss, A.S., Gao, X., Morgan, M.A., Cook, M., Smith, E.R., and Shilatifard, A. (2013). The Mll2 branch of the COMPASS family regulates bivalent promoters in mouse embryonic stem cells. Nat Struct Mol Biol 20, 1093–1097.

Jenuwein, T., and Allis, C.D. (2001). Translating the histone code. Science 293, 1074–1080.

Konze, K.D., Ma, A., Li, F., Barsyte-Lovejoy, D., Parton, T., Macnevin, C.J., Liu, F., Gao, C., Huang, X.P., Kuznetsova, E., et al. (2013). An orally bioavailable chemical probe of the Lysine Methyltransferases EZH2 and EZH1. ACS Chem Biol 8, 1324–1334.

Kriaucionis, S., and Heintz, N. (2009). The nuclear DNA base 5-hydroxymethylcytosine is present in Purkinje neurons and the brain. Science 324, 929–930.

Kumar, D., Cinghu, S., Oldfield, A.J., Yang, P., and Jothi, R. (2021). Decoding the function of bivalent chromatin in development and cancer. Genome Res 31, 2170–2184.

Lawrence, M., Gentleman, R., and Carey, V. (2009). rtracklayer: an R package for interfacing with genome browsers. Bioinformatics 25, 1841–1842.

Liao, Y., Smyth, G.K., and Shi, W. (2019). The R package Rsubread is easier, faster, cheaper and better for alignment and quantification of RNA sequencing reads. Nucleic Acids Res 47, e47.

Love, M.I., Hogenesch, J.B., and Irizarry, R.A. (2016). Modeling of RNA-seq fragment sequence bias reduces systematic errors in transcript abundance estimation. Nat Biotechnol 34, 1287–1291.

Mellen, M., Ayata, P., and Heintz, N. (2017). 5-hydroxymethylcytosine accumulation in postmitotic neurons results in functional demethylation of expressed genes. Proc Natl Acad Sci U S A 114, E7812–E7821.

Mikkelsen, T.S., Ku, M., Jaffe, D.B., Issac, B., Lieberman, E., Giannoukos, G., Alvarez, P., Brockman, W., Kim, T.K., Koche, R.P., et al. (2007). Genome-wide maps of chromatin state in pluripotent and lineage-committed cells. Nature 448, 553–560.

Mohn, F., Weber, M., Rebhan, M., Roloff, T.C., Richter, J., Stadler, M.B., Bibel, M., and Schubeler, D. (2008). Lineage-specific polycomb targets and de novo DNA methylation define restriction and potential of neuronal progenitors. Mol Cell 30, 755–766.

Neri, F., Incarnato, D., Krepelova, A., Rapelli, S., Pagnani, A., Zecchina, R., Parlato, C., and Oliviero, S. (2013). Genome-wide analysis identifies a functional association of Tet1 and Polycomb repressive complex 2 in mouse embryonic stem cells. Genome Biol 14, R91.

Ozawa, P.M., Ariza, C.B., Ishibashi, C.M., Fujita, T.C., Banin-Hirata, B.K., Oda, J.M., and Watanabe, M.A. (2016). Role of CXCL12 and CXCR4 in normal cerebellar development and medulloblastoma. Int J Cancer 138, 10–13.

Patro, R., Duggal, G., Love, M.I., Irizarry, R.A., and Kingsford, C. (2017). Salmon provides fast and bias-aware quantification of transcript expression. Nat Methods 14, 417–419.

Piunti, A., and Shilatifard, A. (2016). Epigenetic balance of gene expression by Polycomb and COMPASS families. Science 352, aad9780.

Rudenko, A., Dawlaty, M.M., Seo, J., Cheng, A.W., Meng, J., Le, T., Faull, K.F., Jaenisch, R., and Tsai, L.H. (2013). Tet1 is critical for neuronal activity-regulated gene expression and memory extinction. Neuron 79, 1109–1122.

Santos-Rosa, H., Schneider, R., Bannister, A.J., Sherriff, J., Bernstein, B.E., Emre, N.C., Schreiber, S.L., Mellor, J., and Kouzarides, T. (2002). Active genes are tri-methylated at K4 of histone H3. Nature 419, 407–411.

Saunders, A., Macosko, E.Z., Wysoker, A., Goldman, M., Krienen, F.M., de Rivera, H., Bien, E., Baum, M., Bortolin, L., Wang, S., et al. (2018). Molecular Diversity and Specializations among the Cells of the Adult Mouse Brain. Cell 174, 1015–1030 e1016.

Serrano, M. (2018). Epigenetic cerebellar diseases. Handb Clin Neurol 155, 227–244.

Singh, S., Howell, D., Trivedi, N., Kessler, K., Ong, T., Rosmaninho, P., Raposo, A.A., Robinson, G., Roussel, M.F., Castro, D.S., et al. (2016). Zeb1 controls neuron differentiation and germinal zone exit by a mesenchymal-epithelial-like transition. Elife 5.

Solecki, D.J., Model, L., Gaetz, J., Kapoor, T.M., and Hatten, M.E. (2004). Par6alpha signaling controls glial-guided neuronal migration. Nat Neurosci 7, 1195–1203.

Solecki, D.J., Trivedi, N., Govek, E.E., Kerekes, R.A., Gleason, S.S., and Hatten, M.E. (2009). Myosin II motors and F-actin dynamics drive the coordinated movement of the centrosome and soma during CNS glial-guided neuronal migration. Neuron 63, 63–80.

Stoyanova, E., Riad, M., Rao, A., and Heintz, N. (2021). 5-Hydroxymethylcytosine-mediated active demethylation is required for mammalian neuronal differentiation and function. Elife 10.

Strahl, B.D., and Allis, C.D. (2000). The language of covalent histone modifications. Nature 403, 41–45.

Tomoda, T., Bhatt, R.S., Kuroyanagi, H., Shirasawa, T., and Hatten, M.E. (1999). A mouse serine/threonine kinase homologous to C. elegans UNC51 functions in parallel fiber formation of cerebellar granule neurons. Neuron 24, 833–846.

van der Heijden, M.E., and Sillitoe, R.V. (2021). Interactions Between Purkinje Cells and Granule Cells Coordinate the Development of Functional Cerebellar Circuits. Neuroscience 462, 4–21.

Verma, N., Pan, H., Dore, L.C., Shukla, A., Li, Q.V., Pelham-Webb, B., Teijeiro, V., Gonzalez, F., Krivtsov, A., Chang, C.J., et al. (2018). TET proteins safeguard bivalent promoters from de novo methylation in human embryonic stem cells. Nat Genet 50, 83–95.

Villasenor, R., Pfaendler, R., Ambrosi, C., Butz, S., Giuliani, S., Bryan, E., Sheahan, T.W., Gable, A.L., Schmolka, N., Manzo, M., et al. (2020). ChromID identifies the protein interactome at chromatin marks. Nat Biotechnol 38, 728–736.

Vladoiu, M.C., El-Hamamy, I., Donovan, L.K., Farooq, H., Holgado, B.L., Sundaravadanam, Y., Ramaswamy, V., Hendrikse, L.D., Kumar, S., Mack, S.C., et al. (2019). Childhood cerebellar tumours mirror conserved fetal transcriptional programs. Nature 572, 67–73.

Watt, A.J., Cuntz, H., Mori, M., Nusser, Z., Sjostrom, P.J., and Hausser, M. (2009). Traveling waves in developing cerebellar cortex mediated by asymmetrical Purkinje cell connectivity. Nat Neurosci 12, 463–473.

Wickham, H. (2016). ggplot2: Elegant Graphics for Data Analysis. Springer-Verlag New York.

Wickham, H., Averick, M., Bryan, J., Chang, W., McGowan, L.D., François, R., Grolemund, G., Hayes, A., Henry, L., Hester, J., et al. (2019). Welcome to the tidyverse. Journal of Open Source Software 4, 1686.

Xiang, Y., Zhang, Y., Xu, Q., Zhou, C., Liu, B., Du, Z., Zhang, K., Zhang, B., Wang, X., Gayen, S., et al. (2020). Epigenomic analysis of gastrulation identifies a unique chromatin state for primed pluripotency. Nat Genet 52, 95–105.

Xu, X., Stoyanova, E.I., Lemiesz, A.E., Xing, J., Mash, D.C., and Heintz, N. (2018). Species and cell-type properties of classically defined human and rodent neurons and glia. Elife 7.

Yu, G., Wang, L.G., Han, Y., and He, Q.Y. (2012). clusterProfiler: an R package for comparing biological themes among gene clusters. OMICS 16, 284–287.

Yu, G., Wang, L.G., and He, Q.Y. (2015). ChIPseeker: an R/Bioconductor package for ChIP peak annotation, comparison and visualization. Bioinformatics 31, 2382–2383.

Zhang, J., Ji, F., Liu, Y., Lei, X., Li, H., Ji, G., Yuan, Z., and Jiao, J. (2014). Ezh2 regulates adult hippocampal neurogenesis and memory. J Neurosci 34, 5184–5199.

Zhu, X., Girardo, D., Govek, E.E., John, K., Mellen, M., Tamayo, P., Mesirov, J.P., and Hatten, M.E. (2016). Role of Tet1 /3 Genes and Chromatin Remodeling Genes in Cerebellar Circuit Formation. Neuron 89, 100–112.

