## Supplementary Figures for "Histone bivalency regulates the timing of cerebellar granule cell development"

SUPPLEMENTARY FIGURE S1

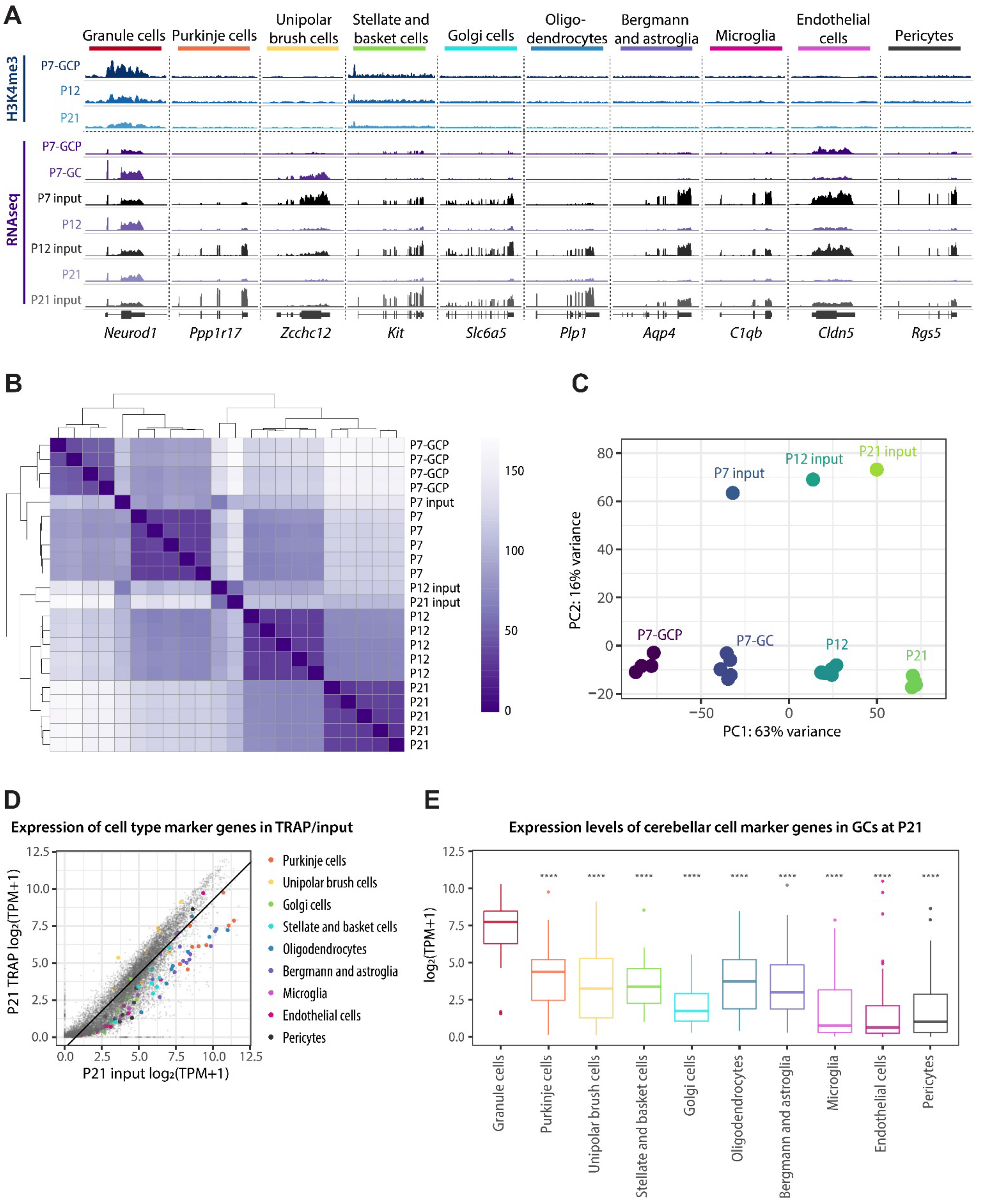


**Supplementary Figure S1**, related to Figure 1. Isolation of GC-specific chromatin and RNA from the postnatal cerebellum. **A** Genome browser view of representative RNA-seq and H3K4me3 ChIPseq signal at marker genes of cerebellar cell types. A representative gene is shown for each major cerebellar cell type, identified using scRNA-seq data from the adult cerebellum (Saunders et al. 2018). Representative ChIP-seq and TRAP RNA-seq samples are shown. H3K4me3 (n=2-3 samples/group) and H3K27me3 (n=4-5 samples/group) ChIP-seq was performed on chromatin isolated from GCPs (P7) or from GC nuclei sorted using FANS (P12 and P21). RNA-seq was performed on TRAP RNA isolated from P5-P7 *Tg(Atoh1-Egfp-L10a)* GCPs (P7-GCP, n=4 samples/group), or from the cerebellar lysates of *Tg(Neurod1-Egfp-L10a)* mice at P7 (P7-GC), P12 and P21 (n=5 mice/group). For input RNA-seq, individual input samples at each age were pooled in equal amounts to yield one input per developmental time point. **B** TRAP RNA-seq sample similarity matrix. **C** Principal component analysis on TRAP RNAseq samples. **D** Scatterplot showing the expression of marker genes of cerebellar cells in P21 TRAP samples relative to input. The top 10 most highly specific genes for each cell type are shown. The line denotes the cut-off level for identifying genes depleted in TRAP samples relative to input. **E** The expression of cell type marker genes in P21 GCs. One-way ANOVA, followed by Tukey HSD *post hoc* test. Correction for multiple comparisons was performed considering all comparisons but significances are shown only for comparisons with granule cells. **** $p < 0.0001$.

SUPPLEMENTARY FIGURE S2

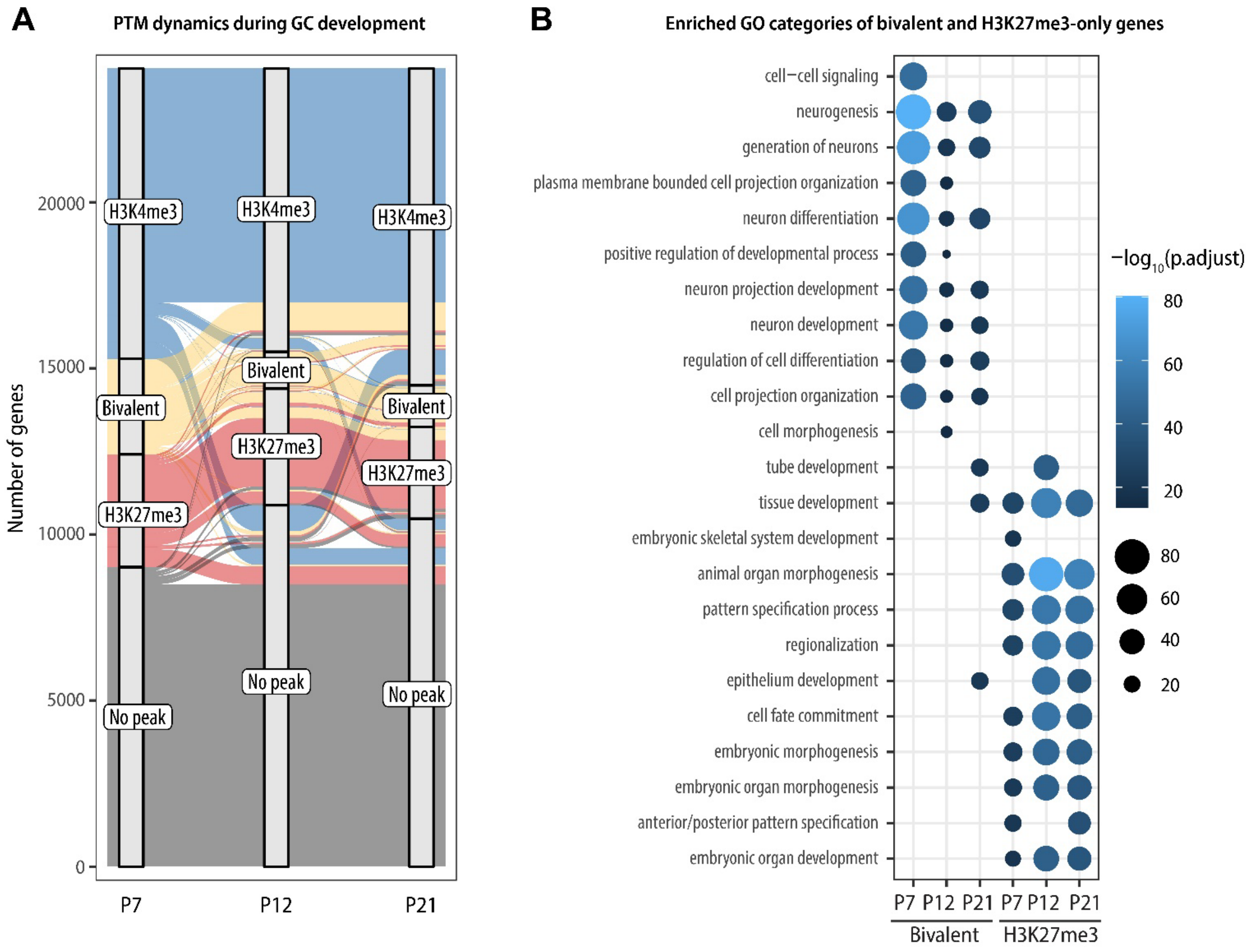


**Supplementary Figure S2**, related to Figures 2 and 4. **A** Alluvial plot showing the dynamics of promoter-proximal histone PTMs between P7 and P21. Groups that contain fewer than 0.5% of included genes are omitted for simplicity. The majority of H3K4me3-only and no-peak genes are stable throughout development, whereas bivalent and H3K27me3-only genes are more likely to change their PTM status. **B** Enriched GO (biological process) categories of genes with bivalent or H3K27me3-only promoters at key stages of GC development. GO categories were identified using clusterProfiler. The GO Biological Process categories were sorted by the adjusted P-value and the top 10 enriched categories are shown for each age.

SUPPLEMENTARY FIGURE S3

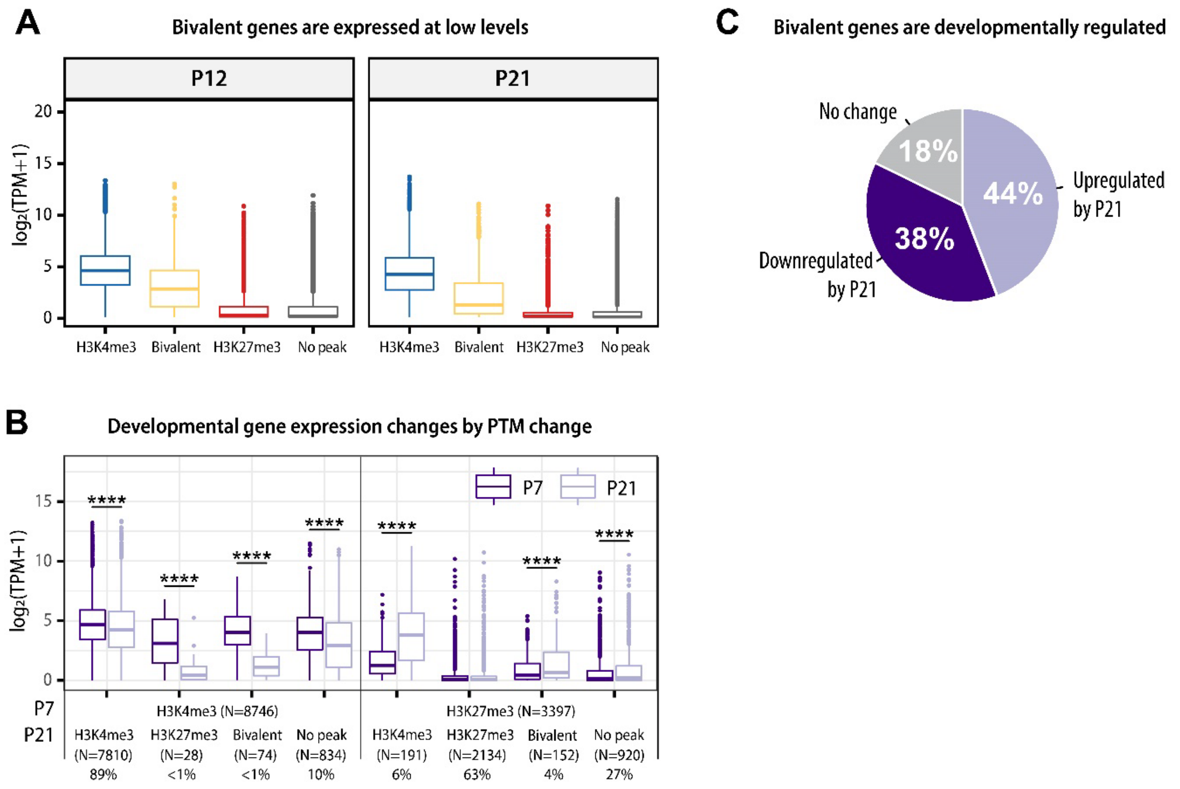


**Supplementary Figure S3**, related to Figure 4. Changes in bivalency correlate with gene expression. **A** The expression of H3K4me3-only, bivalent, H3K27me3-only, and no-peak genes at P12 and P21. **B** Developmental changes in PTM status at the TSS are associated with developmental gene expression changes between P7 GCPs and P21 GCs. P7 H3K4me3-only and P7 H3K27me3-only genes are shown. Pairwise t-test, adjusted for multiple comparisons using the BH method. ** $p < 0.01$, **** $p < 0.0001$. **C** Pie chart depicting the percentage of bivalent genes that are upregulated, downregulated, or do not change in expression between P7 and P21.

SUPPLEMENTARY FIGURE S4

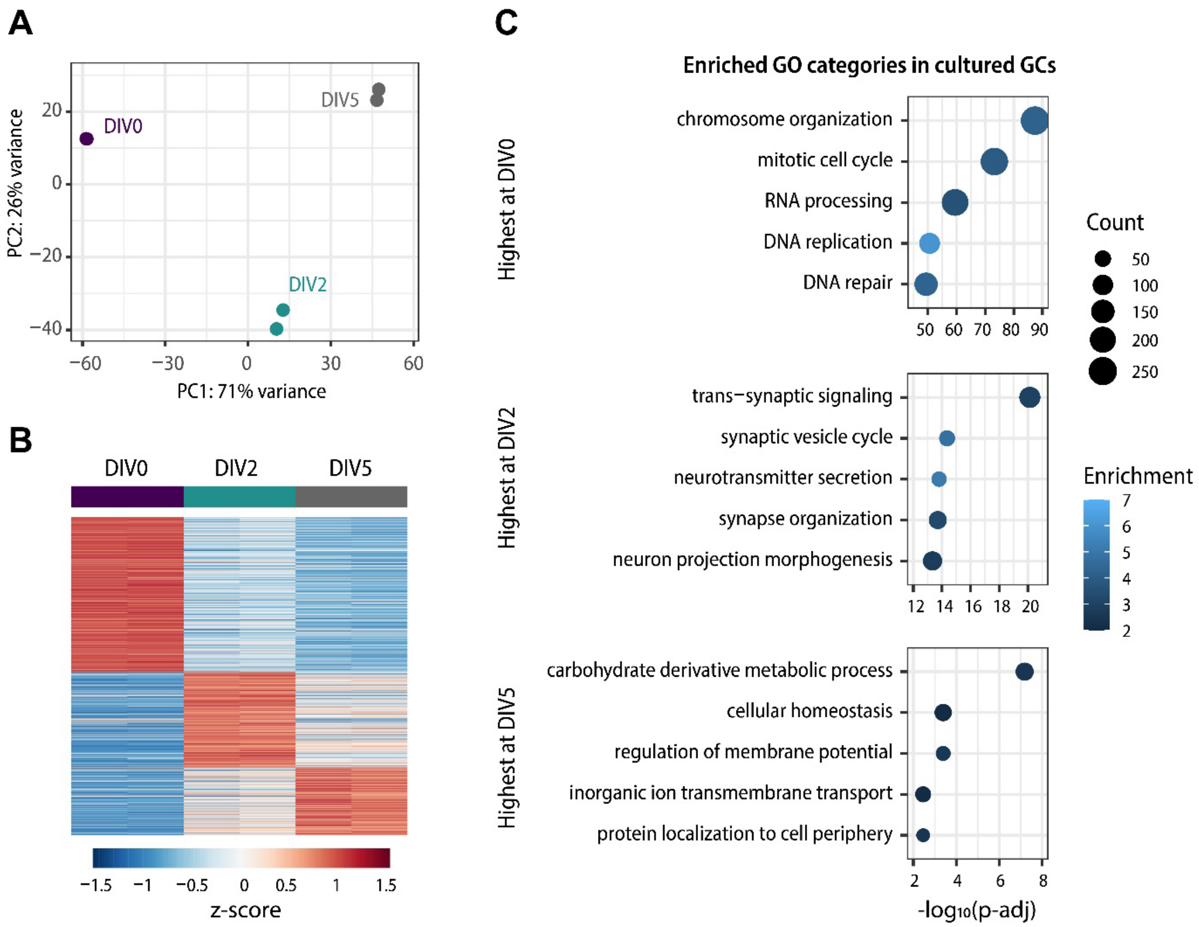


**Supplementary Figure S4**, related to Figure 6. Cultured GCs follow *in vivo* GC developmental trajectories. **A** Principal component analysis on RNA-seq samples isolated from cultured GCs at DIV0, DIV2 and DIV5, cultured in the presence of DMSO. **B** Heatmap depicting differentially expressed genes between cultured GCs at DIV0, DIV2, and DIV5. DE genes (p-adj < 0.05) were identified by pairwise comparisons between groups using DESeq2 and sorted by the highest expressed genes at each age. **C** Gene Ontology analysis of the highest expressed genes in each group. The GO biological process categories were sorted by the adjusted P-value and the top 5 enriched non-redundant categories are shown for each group.

SUPPLEMENTARY FIGURE S5

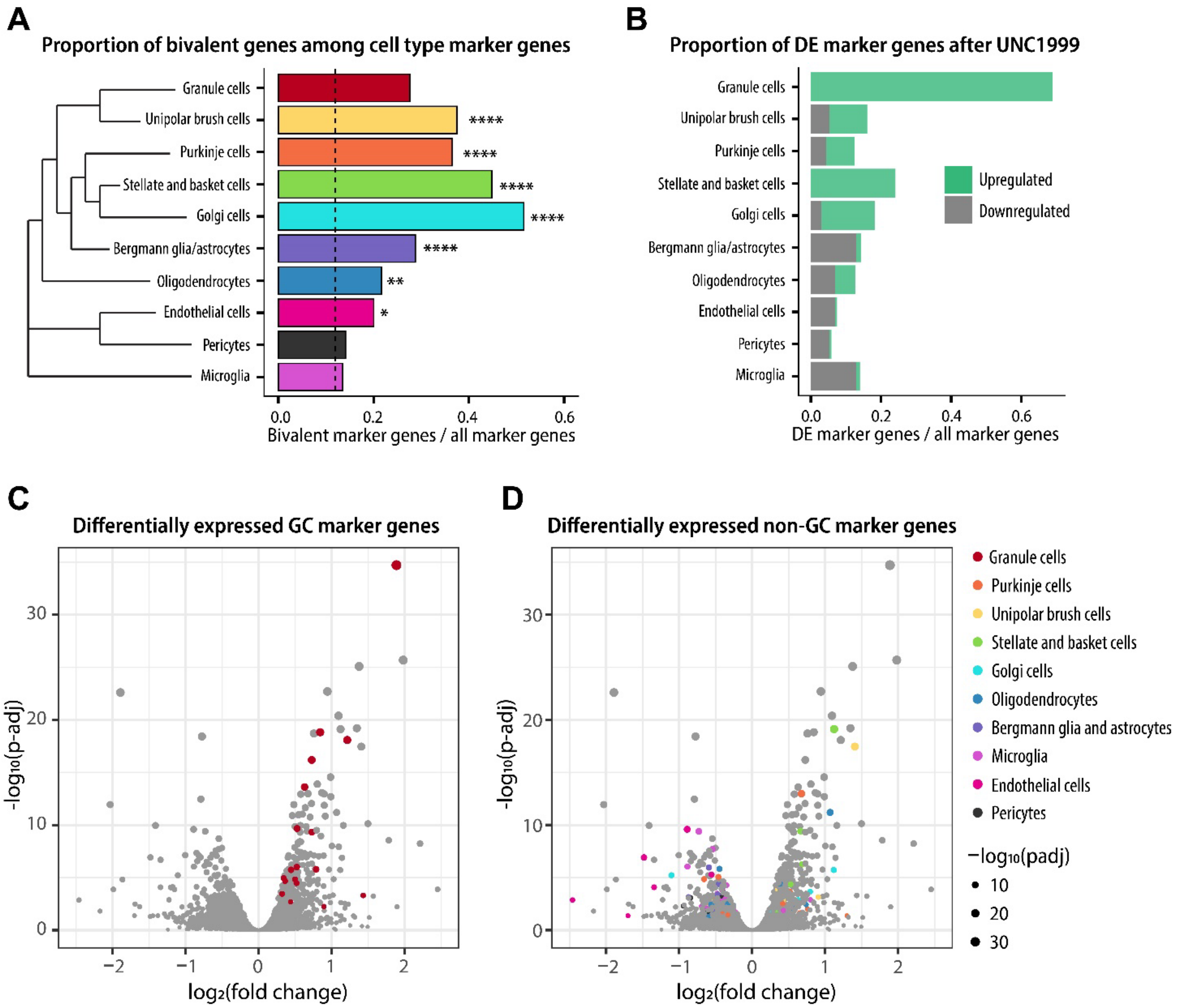


**Supplementary Figure S5**. Regulation of cell type marker gene expression by bivalency in developing GCs. **A** The proportion of bivalent genes among marker genes of cerebellar cells at P7. The genome-wide proportion of bivalent genes is indicated with a dashed line. Pairwise Fisher test was performed between the genome-wide proportion of bivalent genes and the proportion of bivalent genes among each cell type marker genes, using the rstatix package. P-values were adjusted for multiple comparisons using the fdr method. * $p < 0.05$, ** $p < 0.01$, *** $p < 0.001$, **** $p < 0.0001$. **B** The proportion of differentially expressed marker genes of cerebellar cells after UNC1999 treatment. **C-D** Volcano plots showing the differential expression of GC marker genes (**C**, highlighted in red) and marker genes of other cerebellar cells (**D**) in response to UNC1999 treatment.
